# Neural mechanisms of awareness of action

**DOI:** 10.1101/2024.08.15.608153

**Authors:** David S. Jin, Oumayma Agdali, Taruna Yadav, Sharif I. Kronemer, Sydney Kunkler, Shweta Majumder, Maya Khurana, Marie C. McCusker, Ivory Fu, Emily J. Siff, Aya Khalaf, Kate L. Christison-Lagay, Shanae L. Aerts, Qilong Xin, Jing-Jing Li, Sarah H. McGill, Michael J. Crowley, Hal Blumenfeld

## Abstract

The origins of awareness of action (AoA), the ability to report an action just performed, remain elusive. Differing theories ascribe AoA to pre-action, efferent motor/volitional mechanisms versus post-action, afferent sensory/perceptual neural mechanisms. To study these two types of mechanisms and others, we developed a paradigm where very similar aware and unaware actions occur repeatedly. Aware actions demonstrated larger neurophysiological signals both preceding and following movement. The differences included well-known volitional and perceptual event related potentials (PMP, N140, P300), as well as frontal midline theta, event-related alpha/beta desynchronization, and post-move blink rates. On longer time scales, we identified a novel event related potential preceding unaware moves, and found behavioral and pupillometric evidence for decreased attention and arousal over minutes concurrent with AoA loss. Our findings suggest that both dynamic, individual action-associated volitional and perceptual neural activity, as well as long-term attention and arousal states play a role in maintaining AoA.

## Introduction

The neural origins of awareness of action (AoA), the ability to report an action just performed, have long been poorly understood, yet have been debated from the inception of psychological science. The loss of AoA is a part of everyday experience, seen in phenomena like “highway hypnosis” and “white line fever” where awareness of recent driving actions are lost, or being unable to know whether everyday tasks have been completed such as locking the door or turning off the gas[1, 2]. Aberrant AoA is regularly seen in medical conditions such as Parkinson’s disease, ataxia, and schizophrenia[3, 4]. Despite the lack of understanding of neural mechanisms of AoA, definitions of AoA are also considered in legal judgments, with distinctions between murder versus manslaughter and voluntary versus involuntary crimes, depending on whether the offenses were committed with intention and awareness[5-7].

The early founders of modern psychological science fell into two primary camps on the origins of awareness of motor action, creating the “Two Williams Debate”. The first camp, led by Wilhelm Wundt, posited that the contents of action are represented *a priori*, and that AoA is a mental, generative phenomenon independent of somatosensory feedback. The second camp, led by William James, suggested that the contents of action are represented *a posteriori*, and that the somatosensory, afferent processes allow for AoA[8]. We posit that neither perspective fully encapsulates the neural processes underlying AoA. Volitional as well as perceptual neural mechanisms both pre- and post-action may be needed to establish AoA on short time scales, while additional state-related attention and arousal mechanisms could influence AoA over longer time periods.

To test whether AoA is an *a priori* or *a posteriori* process (Two Williams Debate), a carefully designed, behavioral paradigm to isolate instances of awareness or unawareness is needed. While previous studies of action have included varying motor components such as button presses when the subject feels the urge to move (e.g. the classic Libet paradigm), forced choices, and even bungee jumping, they do not query whether the subjects are aware of the identity of the action or not[9-11]. Therefore, they have not tested AoA itself, by looking for neural signals seen when AoA is present versus absent in a controlled setting. To address this, we developed a paradigm based upon a classic sliding block puzzle game, which subjects complete at a self-set pace. Periodically, subjects are asked questions about their just-completed move, to assess awareness of the previous action. This approach, referred to as contrastive analysis, where neural signals are compared between very similar events with versus without conscious awareness, has been applied widely to sensory paradigms, but has not been used so far to study motor awareness [12-17]. The paradigm we developed, involving repeated relatively similar clicks and keypresses, allows us to perform a contrastive method to AoA and to observe both pre-action and post-action neural signals.

Should the neural mechanisms of AoA differ *a priori*, differences in neural activity are expected during the preparation and initiation of action. Previous studies of action, including the classic Libet paradigm, have established pre-action differences in awareness of intent to act[18, 19]. These studies have established the presence of the *bereitschaftspotential* (BP), or readiness potential, a negativity in EEG 1-2 seconds preceding the intent to act[10, 18-21]. The readiness potential has been shown to be generated by activity in supplementary motor cortex and premotor cortex[22-25]. Much closer to performance of the action (approximately 100 ms prior), a pre-movement positivity (PMP) has been observed as well, associated with the immediate initiation of the action as opposed to its planning[26-28]. Additionally, pre-action activation changes in prefrontal and parietal cortex have been shown to occur prior to intent to act, seen in both PET and fMRI [29-31]. On the other hand, *a posteriori* differences in neural activity would manifest in perceptual signals, like those seen in the visual, auditory, and tactile domains. Known event-related potentials following perceptual stimuli include the face-specific N170, N100 (vision), auditory awareness negativity, and N140 (somatosensory awareness negativity)[12, 32, 33]. In addition, signals which support both *a priori* volitional and *a posteriori* perceptual hypotheses of AoA are known to exist. In the time-frequency domain, pre-action beta frequency (12-30 Hz) suppression occurs prior to the initiation or imagination of a voluntary movement[34-36], and directives to attend to a visual stimulus are known to decrease the presence of beta activity[37]. Lastly, suppression of alpha activity (8-12 Hz) has long been known to increase the probability of detecting a visual stimulus, and, like beta activity, is also quieted by the presentation of a visual stimulus[38-40] and is quieted prior to the execution of voluntary movement[41]. Alpha suppression is also observed in auditory and tactile domains[42, 43], and alpha/beta event-related desynchronization more generally is viewed as signature of prominent behavioral or perceptual events and conscious awareness[44-46].

The view of action as purely *a priori* or *a posteriori* may also be incomplete. Actions in the real world are rarely performed as individual discretized motions, a drawback of many traditional forced choice and Libet-style studies. Instead, they are completed in sequence and under varying states of arousal. Studies of attentional states suggest that these flow states are bimodal in nature (i.e. “in the zone” or “out of the zone”), modulated by the default mode network, and can be last on the order of seconds to minutes[47-49]. We hypothesized that AoA may be lost more often during mind-wandering and daydreaming circumstances, generally states of lower-arousal or distraction from the task at hand. We encouraged these circumstances by having subjects play the game while an engaging video appeared in the background, which they had to describe following each run. To measure the effects of fatigue and arousal-related factors on awareness over the course of the study, we used pupillometric measurements. Pupil diameter is a known to track with neural activity in the salience network, and can provide rich, noninvasive information on arousal levels[50, 51]. The success of multiple continuous performance task types can be predicted by pupil diameter[52]. Decreased arousal in tandem with decreased AoA would provide concrete evidence for this anecdotal link.

We found clear differences in event-related potential, EEG time-frequency, and eye metric data both before and after action. We also found that arousal levels, measured by pupil diameter, decreased throughout the course of the experiment, in tandem with decreased levels of behavioral awareness and increased levels of unawareness. Thus, our findings support both hypotheses of the Two Williams Debate: that generative, volitional processes and afferent, perceptive processes allow for AoA. Our findings also highlight the role of longer-term attention and arousal states in the maintenance of AoA and that AoA must be considered in a broader context of sequences of actions.

## Methods

### Clinically Healthy Participants

Sixty-seven clinically normal, adult subjects were recruited for our study. Inclusion criteria included normal vision (with soft lens correction) and normal hearing. Exclusion criteria included current or past diagnoses of neurological disorders and vision correction which required hard lenses. All subjects underwent EEG and pupillometry measurements. Summary data for healthy participants are shown in Table 1.

### Behavioral Exclusionary Criteria

A total of 10 subjects were excluded from the EEG sample due if their total number of aware trials and/or unaware trials fell below a 12 trial minimum following individual trial rejection (See hdEEG – Data Preprocessing; Pupillometry – Individual Trial Rejection). Individual trials were labeled as having excessive artifact labeled by our automated pipeline. A similar 12-trial minimum was applied to the pupillometry sample and a total of 11 subjects were excluded due to tracker loss or noise. Thus, our sample included 57 EEG subject datasets and 56 pupillometry subject datasets. For our analyses of video engagement and familiarity, 46 datasets were collected. Of these subjects, one had previously seen the videos as a participant in a pilot version of the task. This subject was therefore excluded from analyses of video familiarity. Additionally, block identity data for 12 of 67 subjects were not saved due to a coding error; thus, our block identity chi-squared analyses were based on 55 of 67 subjects.

## Behavioral Task

### Equipment and Hardware

Subjects completed the task using a gameplay mouse (Lenovo Essential USB Mouse) and keyboard (Lenovo Essential Wired Keyboard) attached via USB to the task laptop (Figure S1). In addition, for a free recall session, subjects spoke into a microphone (Snowball iCE, Logitech). Subjects viewed the game on a 17-inch LCD monitor an arm mount (EyeLink 1000 Plus System, SR Research, Inc.) displaying the task laptop (MSI Gf63 Thin, MSI) visual feed. The LCD screen was connected to the experiment laptop via a VGA/HDMI adapter. The task code was implemented in the *PsychoPy* package of Python.

### Task Progression

Subjects performed the task across three days, with one training day and two testing days (**Fig 1. A**). Six ten-minute runs were administered on each day for a total of one hour of gametime (**Fig 1B**). To complete the task, subjects had to navigate a red block out of the bounds of a grid, by moving obstructing blocks to free a path for the block. To move a block, subjects used the mouse in the right hand to click on and select a block. They then used the WASD keys in the left hand (W = up, middle finger; A = left, ring finger; S = down, middle finger; D = right, index finger) to move a block. Upon completion of the movement, subjects pressed the spacebar with the left thumb to confirm the action. Subjects could not select and move another block without first confirming the current block movement. Event related potential, time-frequency, and pupillometry analyses were time-locked to the confirmation of the action, and all further uses of “time from action” refer to timeframe with respect to the confirmation. When the subject solved an individual puzzle, a message depicting “Good Job” would appear on the right third of the screen, after which the next puzzle configuration would be shown. A randomized rotation of ten unique puzzle configurations was used. Every 2-5 moves, the board disappeared for a period of 2-8 seconds before reappearing. Subjects were trained on this task for three runs with a plain blue background.

**Figure 1.**
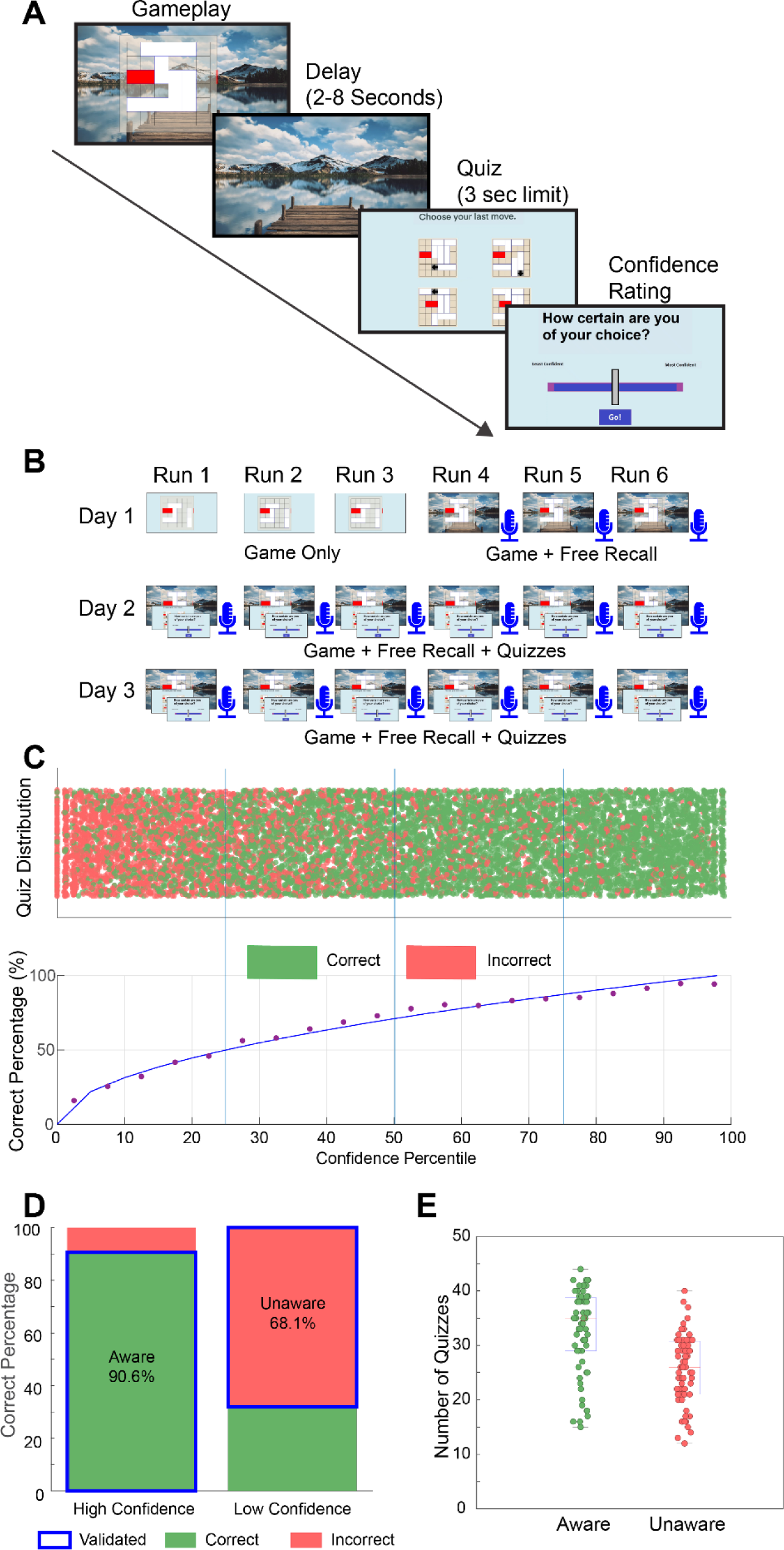
A. Experimental task. Subjects (N = 67) played the task game by navigating a red block out of the bounds of the grid by moving obstructing blocks out of the way. Periodically, the game board would disappear, and a two part quiz could be shown: 1. A multiple choice question in which subjects selected the move just performed (direction and block) and 2. Indicated their confidence in their choice. B. Experimental progression. The experiment consisted of one training and two testing days. On the training day, subjects were first trained on a fixed set of puzzle configurations for three 10-minute run. Next, subjects completed the puzzles while a background distractor video played. After each run, subjects had to recount what they recalled from the distractor video. Subjects completed three runs with the background distractor on the training day. The two testing days were identical. Subjects completed six runs with the background distractor and with quizzes for awareness administered periodically. C. All quiz results amongst all subjects (n = 9,966). Any quiz in which the subject correctly identified the block and indicated high confidence was designated aware. Any quiz in which the subject incorrectly identified the block and indicated low confidence was designated unaware. High confidence was designated as any confidence above the 75^th^ percentile of the subject’s answers. Low confidence was designated as any confidence below the 25^th^ percentile of the subjects answers. D. Validation rates. Amongst high confidence responses, 90.6% were correct (expected: 100% correct), and amongst low confidence responses, 68.1% were incorrect (expected: 75% incorrect as the subject had a ¼ chance of randomly guessing correctly). E. Subject aware and unaware trial totals. Subjects attained an average of 32.6 aware and 25.5 unaware trials (both testing day totals summed).

In latter three runs of the training day, subjects performed the task with a distractor task added, in which they were instructed that at the end of each run, they would have to freely recall as many details as possible in a background video that played as they performed the task. Subjects were instructed to maintain a chosen pace of play even with the background video playing. At the end of each run, subjects completed a three-minute recall session in which they were instructed to continue recounting details until the end of the three minute period. Subjects completed three runs of this task. The two testing days were identical to one another. On the testing days, the game with video background and free recall session was administered. However, after the board disappeared following 2-5 moves, there was a 50% probability that a two-part quiz would be shown. The first part of the quiz was a four-part multiple choice question, in which four candidate moves (with the block and direction indicated) were presented to the subject (with the instruction to “Choose your last move.”). Subjects clicked on which of the four moves they thought was their previous move. The second part of a quiz was a confidence indication. Subjects were asked to indicate their confidence level (“How certain are you of your choice?”) by clicking on a slider (leftmost = least confident, rightmost = most confident). Subjects then pressed a “Go” key to submit their answer, and the game with video resumed.

To discourage subjects from trying to recalculate a particular route taken in the game, as opposed to immediately demonstrating awareness of the previous action, a three-second time limit was imposed for the multiple choice question. If subjects did not answer the multiple choice question within the three second limit, the confidence question was not administered. Late answers were excluded from analysis due to the possibility of mouse slips. During the first three runs, subjects were monitored for excessive numbers of late quizzes. After the first three runs of the experiment, subjects’ late trials were tabulated. If a subject averaged at least two late quizzes per run, or had three or more late quizzes in a single run, subjects were instructed, “Please provide an answer within the three second limit”.

### Data Processing

After each run, five types of behavioral data were saved: 1. TTL pulse timing, 2. Multiple choice accuracy, 3. Confidence slider positions, 3. Order of puzzles presented to the subject, 4. Initial configurations prior to a move being performed, and 5. Background movie identity. These were saved in a *.psydat* file, constituting the behavioral data files from which awareness was determined.

## Pupillometry

### Equipment and Software

Pupillometry and eye metric data were collected using the EyeLink 1000 Plus System (v5.09, SR Research) running on a Dell desktop PC (Model D13M; Dell, Inc.). Data were sampled at 1000Hz using a 35 mm camera and an infrared illuminator mounted below the LCD game monitor (**Figure S1**). Prior to the first and fourth runs of the procedure, participant gaze position was calibrated using a 9-point visual gaze sequence. Additionally, prior to the first run of the experiment, corneal and pupil thresholds for reflectivity were determined. To stabilize head positions, participants performed the task in a chinrest set 55 cm away from the EyeLink camera.

### Data Processing

EyeLink recordings were saved following each run. Raw EyeLink data consisted of two items: 1. Pupil diameter, 2. x/y gaze position, and 3. Ethernet synchronization messages. We extracted three datasets from the raw data: 1. Z-scored pupil diameter, 2. Blink rate, and 3. Saccade rate. Firstly, blinks were identified at the whole run level using the *Stublinks* procedure, which initially downsampled the data of 1000 Hz timepoints to 60 Hz samples. *Stublinks* identifies blinks as changes in pupil diameter fulfilling the following requirements: 1. Diameter changes greater than 0.5 mm between consecutive samples, 2. Diameter differences < 0.1 mm or > 4 mm from the median pupil diameter of the entire run, and 3. Diameter changes 0.4 mm across four consecutive samples, 4. Timepoints with > 1 mm difference between 60 Hz downsampled and 1000 Hz raw timecourses, and 5. Samples with diameter outside of the Tukey’s test IQR. Any period of consecutive artifactual timepoints lasting 100-1000 ms was deemed a blink. A binary vector containing all timepoints was then generated (1 = blink, 0 = non-blink) to represent the run. All artifactual timepoints were linearly interpolated with adjacent non-artifactual samples for creating mean timecourses of pupil diameter. Saccade occurrences in timecourses were determined by identifying timepoints in which gaze position subtended a velocity > 3 degrees/second and < 7 degrees/second of visual angle in a 5 ms duration. As with blink data, a binary vector containing all timepoints was then generated (1 = saccade, 0 = non-saccade) to represent the presence of saccades in the run. Following initial processing of raw data, pupil diameters were z-scored. Given that the luminance of the quiz was brighter than that of gameplay, all timepoints in a run that corresponded to the gameplay only were identified. To account for recalibration after run 3, all non-quiz timepoints of runs 1-3 were aggregated and the mean and standard deviation of all three runs in aggregate was calculated. Individual gameplay pupil diameters were then z-scored according to these mean and standard deviation values. The same process was repeated for runs 4-6. We then extracted 8-second epochs (4 seconds prior to 4 seconds following) centered around the confirmation of an action immediately preceding the board disappearance, leading up to a quiz. These trial epochs were identified with Ethernet timing messages communicated from the behavioral laptop, and were extracted for raw pupil data, z-scored pupil data, blink rates, and saccade rates.

### Individual Trial Rejection

To identify tracker loss, a period 2 seconds before to 2 seconds following action was isolated within each individual trial. The starting pupil diameter of the isolated period was determined and the end point pupil diameter determined. A 4000-timepoint vector was then generated representing a straight line between the pupil’s starting diameter and the end diameter. The Pearson correlation between this vector and the actual −2 to 2 second period of data were computed. Any trial with a r-value greater than 0.99 was eliminated, as the only circumstance under which the actual pupil trace would resemble a straight line would be if the tracker was lost. In these cases, *Stublinks* would isolate the point before the loss of the tracker and the point after the tracker began tracking once again, and interpolate a straight line between these. Such a high r-value could also occur if the tracker had tracked a miscellaneous object of fixed diameter (e.g. an EEG electrode).

## hdEEG

### Equipment and Software

EEG data were collected with 257-lead Ag/AgCl electrode nets (Hydrocel GSN 256, Magstim EGI Inc), sampled at 1000 Hz. Two systems were used: 1. Two Net Amps 200 128-Channel Amplifiers, and 2. Net Amps 400 256-Channel Amplifier. Recordings were made on a 1. desktop computer (Power Mac G5 Quad; Mac OS X v10.5.8, Apple, Inc.) running NetStation 4.2.2 and 2. (A Macbook Pro 2018; MacOsX 10.14.2). Signals were acquired as Cz-referenced.

### Data Preprocessing

EEG data were preprocessed with the EEGLAB (function name in parenthesis) at the session and the individual epoch level. Session level processing began with an initial 1 Hz high-pass filter First, data were high-pass filtered (> 1Hz) to correct for drift (function *clean drifts* with parameters for 0.25 and 0.75 Hz). Next, 60 Hz line noise was removed from the data (function *pop_cleanline*, with parameters for 60 and 120 Hz). Following cleaning of line noise, remaining noisy channels were rejected (function *clean_channels*, with parameters 0.8, 0.5, and 4) and noisy data timepoints were found (function *clean_windows* with parameters -Inf, 7, and 0.25). The data were then re-interpolated with spherical interpolation (function *pop_interp*). Data were then re-referenced to the common average reference.

At the epoch level, epochs of 4000 ms (2000 ms prior to action and 2000 ms following) were isolated from the data based upon timing synchronization flags. These trials were then rejected if the quantity of noisy timepoints between 200 ms pre-action and 500 ms following action exceeded 175 ms (i.e. 25% of timepoints). The kept trials were then concatenated into a single vector (*channels* x *timepoints* x *epochs*). This vector was passed through a 10-component principal component analysis (PCA), then through independent component analysis (ICA) upon the PCA data. Components corresponding to blink, saccade, cardiac, and myographic artifacts were identified by eye and rejected, and the remaining components recomposed to form the final subject data.

Time-frequency maps were extracted using wavelet decomposition (continuous wavelet transform; CWT) and individual epoch rejection was performed with short-time Fourier transform (STFT). Wavelet transformation was chosen as it demonstrated superiority in maintaining both temporal precision while also accounting for the disparity in duration of slow versus fast waves[53]. STFT was chosen for epoch rejection as high-power, artifactual frequencies would require maintenance over longer periods to be deemed a contaminant. For these data, 6000 ms data vectors (3 seconds pre-action to 3 seconds post-action) were extracted and preprocessed as described above. For the calculation of STFT, individual epochs were first divided into bins of 125 ms duration with 25% overlap. STFT was then performed in MATLAB extracting spectral power for 1-150 Hz (with function *spectrogram*) on each channel. These were then squared to give power values. Next, bins corresponding to timepoints 2 seconds pre-action to 1 second pre-action were identified for baselining and mean and standard deviation of power values calculated. All other timebins were then z-scored using these values. The beta and gamma power z-score for each timebin was then calculated by averaging the z-score value of each constituent frequency of the band (12-30 Hz for beta, 40-140 Hz for gamma). If any timebin in any channel of the epoch contained an average beta power z-score value greater than 100, or average gamma power z-score value greater than 100, the trial was excluded.

For statistical analysis and visualization, CWT was performed on the included epochs. First, a discrete Fast Fourier transform was performed on each individual trial. Next, for each frequency we analyzed (1-125 Hz), a Morelet wavelet was generated and convolved with the data. An inverse discrete Fourier transform was then performed on the data. These data were then converted to power values by squaring. For each epoch, channel, and frequency, timepoints corresponding to the period 2 seconds prior to the action to 1 second prior were identified, and the mean and standard deviation taken. The data timepoints within the epoch and the particular channel were then z-scored according to the corresponding mean and standard deviation channel).

## Timing Synchronization

### hdEEG

Task and behavioral events were transmitted from the experimental laptop to an Arduino Uno R3 board via USB to deliver transistor-transistor logic pulses to the EEG amplifier. For the Net Amps 200 system, the pulse was delivered from Arduino to amplifier with a DB9 cable. For the Net Amps 400 system, the pulse was either delivered via DB9 to a clock box which connected to the amplifier via MRTJ cable, or via DB9 to a HyperGrip adapter. Four events were employed: 1. The answering of any quiz question, or late multiple choice message display; 2. Any visual state change (i.e. board disappearance, puzzle completion, quiz display); 3. Confirmation of the action with the spacebar, and 4. Selection of the block with the mouse. Timing testing with a photodiode was employed as described previously[13]. For the Net Amps 200 system and clock box connection, the photodiode reflex appeared 70 ms after the arrival of the TTL pulse; and for the hypergrip cable, the photodiode reflex appeared 78 ms prior to the TTL pulse. Thus, a 148 ms difference was noted between the two connections. Epochs acquired from each system were extracted from raw data with these offsets applied.

### EyeLink

Behavioral performance and pupillometry were synchronized via Ethernet messages sent between the experimental laptop and EyeLink PC (SR Research, Inc.). These messages included timing data on block selection, block movement, action confirmation, quiz display, and quiz answers. In addition to timing details, Ethernet messages also detailed the block selected, movement direction, quiz accuracy, and confidence slider submission position.

## Statistical Analysis

### Behavioral Data

Following behavioral data acquisition, *.psydat* files were read into MATLAB (R2019, *Mathworks*) to extract raw data from the *.psydat* files. Designations for “aware” and “unaware” actions based upon quiz answers contained in the *.psydat* files were analyzed at the session level. The multiple choice answers (either “Y” for correct, or “N” for incorrect) were aggregated across all six runs. The same was done for raw confidence values (−450 to 450, a 900-pixel span centered on the screen). Confidence values were then sorted by percentile. Any correct answer followed by high confidence (>75^th^ percentile within the subject’s data) was marked “Aware”, and any incorrect answer followed by low confidence (<25^th^ percentile within the subject’s data) was marked “Unaware”. These designations were then used to sort aware and unaware epochs in EyeLink and hdEEG data.

One-sample t-tests testing runs 2-5 pairwise with run 1 were used to compare run-by-run effects (p < 0.05, Benjamini-Hochberg false-discovery rate correction). One-sample t-tests were used to compare differences between days (p < 0.05) in awareness, unawareness, confidence, and accuracy. One sample t-tests were used to test awareness rates for each quiz timepoint (i.e. quiz *n-6* prior to an aware quiz was compared against quiz *n-6* prior to an unaware quiz). The twelve timebins were corrected for significance with the Benjamini-Hochberg procedure. Two-sample t-tests were used to compare differences between sexes (p < 0.05) in awareness, unawareness, confidence, and accuracy. Spearman correlation coefficients were used to compare relationships between video engagement and awareness, unawareness, confidence, and accuracy. Spearman correlation coefficients were also used to compare relationships between video familiarity and awareness, unawareness, confidence, and accuracy. χ^2^ tests were used to compare outcomes for awareness and unawareness based on identity (red vs white): 2. Two-category (aware, unaware); 2. Three-category (aware, unaware, other); 3. Five-category (aware, unaware, mid-high confidence, mid-low confidence, unvalidated). Mid-high trials have confidence percentile >50% and ≤75% (regardless of answer on multiple choice); Mid-low trials have confidence percentile ≥25% and ≤50% (regardless of answer on multiple choice); Unvalidated trials have either incorrect move identification on multiple choice and confidence percentile >75% or correct move identification on multiple choice and confidence percentile <25%.

### hdEEG Spatiotemporal Analyses

To correct for multiple comparisons, we performed spatiotemporal cluster-based permutation analyses on our data[54]. In this analysis, an aggregate null distribution is constructed by randomly permuting our EEG data and identifying significant timepoints with spatially or temporally adjacent to one another, which form spatiotemporal clusters. Initially, an aggregate spatiotemporal null distribution was generated for 5000 permutations. For each iteration, baseline data (2 seconds pre-action to 1 second before action) were shuffled with test period data (1 second pre-action to 2 seconds post-action). Each individual timepoint between aware and unaware epochs was tested for significance with a paired 2-tailed t-test and significance level p < 0.05. Spatial adjacency was determined by a binary matrix, representing each electrode and the electrode number (1-257) of adjacent electrodes. Temporal adjacency was determined if two surrounding timepoints were also deemed significant. Following identification of spatiotemporal clusters, the summed absolute t-value (negative and positive clusters were identified differently) was calculated for all clusters. The cluster with the most positive and most negative t-value was then added to the respective aggregate null distribution. Any cluster within the top 5% of the aggregate null distribution was considered to be statistically significant. Timecourse data were constructed using the findings of the aggregate null distribution for voltage and frequency power. For timepoints within timecourses to be considered significant, their constituent spatiotemporal cluster had to exceed set thresholds of temporal duration and spatial extent (20 ms and 20 electrodes for event-related potential voltage, 200 ms and 10 electrodes for frequency power z-score). Additionally, within individual electrode timecourses, the absolute duration of a series of significant timepoints had to exceed 20 ms to be considered as such.

### EyeLink Temporal Analyses

Given the lack of spatial information within pupillometric data (i.e. pupil diameter, blink rate, and saccade rate), clustering along spatial lines was not possible. Therefore, we performed cluster permutation on temporal lines only. As with spatiotemporal cluster permutation, an aggregate temporal null distribution was initially generated with 5000 iterations. For each iteration, baseline data (the average value of all timepoints 2 seconds pre-action to 1 second before action) were permuted with test period data (1 second pre-action to 2 seconds post-action). The permuted data were then tested with a paired one-sample t-test, and as with spatiotemporal clustering, clusters were identified by temporal adjacency (i.e. statistically significant timepoints occurring in sequence), and negative and positive clusters were considered separately. Within cluster, t-values were summed and the cluster with the most positive and most negative t-value sum was then added to the respective aggregate null distributions. Clusters were then identified on the original unpermuted data with the same method as the permuted data (one sample t-test compared to baseline mean), and then tested against the aggregate null distribution.

### Pupil Run-by-Run Data

Run-by-run calculation of pupil data was performed in a similar manner as for run-by-run behavioral metrics. Within individual aware and unaware epochs, the mean of all pupil diameter values one second prior to action until action performance was taken. These were then averaged within subject and then across subjects. One-sample t-tests were then used to compare runs 2-6 against the pupil diameter of run 1 (p < 0.05, Benjamini-Hochberg false-discovery rate correction).

## Results

### Behavioral Results – Task Validation

To determine if our paradigm had reliably induced the loss of awareness of action in a manner that subjects could self-assess and recognize, we assessed the validation rates and frequencies of awareness and unawareness in our data. Actions in which the subject correctly identified the block and direction of move, and indicated high confidence in their answer was deemed “aware”. Actions in which the subject incorrectly identified the block and direction of move, and indicated low confidence in their answer was deemed “unaware”. Confidence was defined relatively within subject; high confidence was defined as being >75^th^ percentile amongst the subject’s confidence submissions, and low confidence was defined as being <25^th^ percentile amongst the subject’s confidence submissions **(Fig 1C)**. We found that if subjects had indicated high confidence on the confidence question, there was a 90.6% probability they had that they had correctly identified the block they had moved (expected: 100%). We also found that if subjects indicated low confidence on the confidence question, there was a 68.1% probability that they had incorrectly identified the block (expected: 75%; given the four options, being at chance level would indicate that only 25% should be correct; **Fig 1D**). Subjects produced a mean of 32.6 ± 8.0 aware and 25.5 ± 6.5 unaware trials summed across the two testing days (**Fig 1E**).

Lastly, we found no relationship between block identity (red block vs. white block) on awareness/unawareness outcomes (**Table S1**). We found no effects of sex on awareness, unawareness, raw confidence, and accuracy (**Table S2**). We found no effects of age on awareness, raw confidence level, and accuracy, and a statistically significant, albeit small relationship between age and unawareness rates (r = −0.29, p = 0.02; **Fig S2**). Of our 67 subjects, 61 completed both testing days of the experiment. We found no effects of testing day on unawareness, awareness, accuracy, nor confidence. (**Table S3**).

To assess the effects of the background distractor task on accuracy and awareness rates, we had subjects rate each video on a 5-point Likert scale for both their familiarity with the video and its subject matter, and their interest and engagement with the video. We found no effects of familiarity and engagement on awareness, unawareness, accuracy, nor confidence, neither within subjects nor across subjects (**Table S4, S5**).

### Event-Related Potentials – Volitional and Perceptual

To understand the pre-action, anticipatory mechanisms underlying AoA, and the post-action perceptual signals underlying AoA, we performed 256-channel high-density scalp EEG on subjects as they completed the Rush Hour Task/ Immediately prior to the performance of the action, we noted a pre-motion positivity (PMP) over frontal regions, which begins ∼100 ms prior to action and peaks at action performance (**Fig. 2A, B**). The amplitude of the PMP was heightened prior to aware actions as compared to unaware actions. Following the action, we noted the presence of a well-known perceptual signal, the somatosensory awareness potential, or N140. The N140 peaked over right parietal regions (contralateral to the hand executing the action and confirmation) ∼120 ms following the action, and was greater in amplitude for aware actions (**Fig 2A, B**).

**Figure 2.**
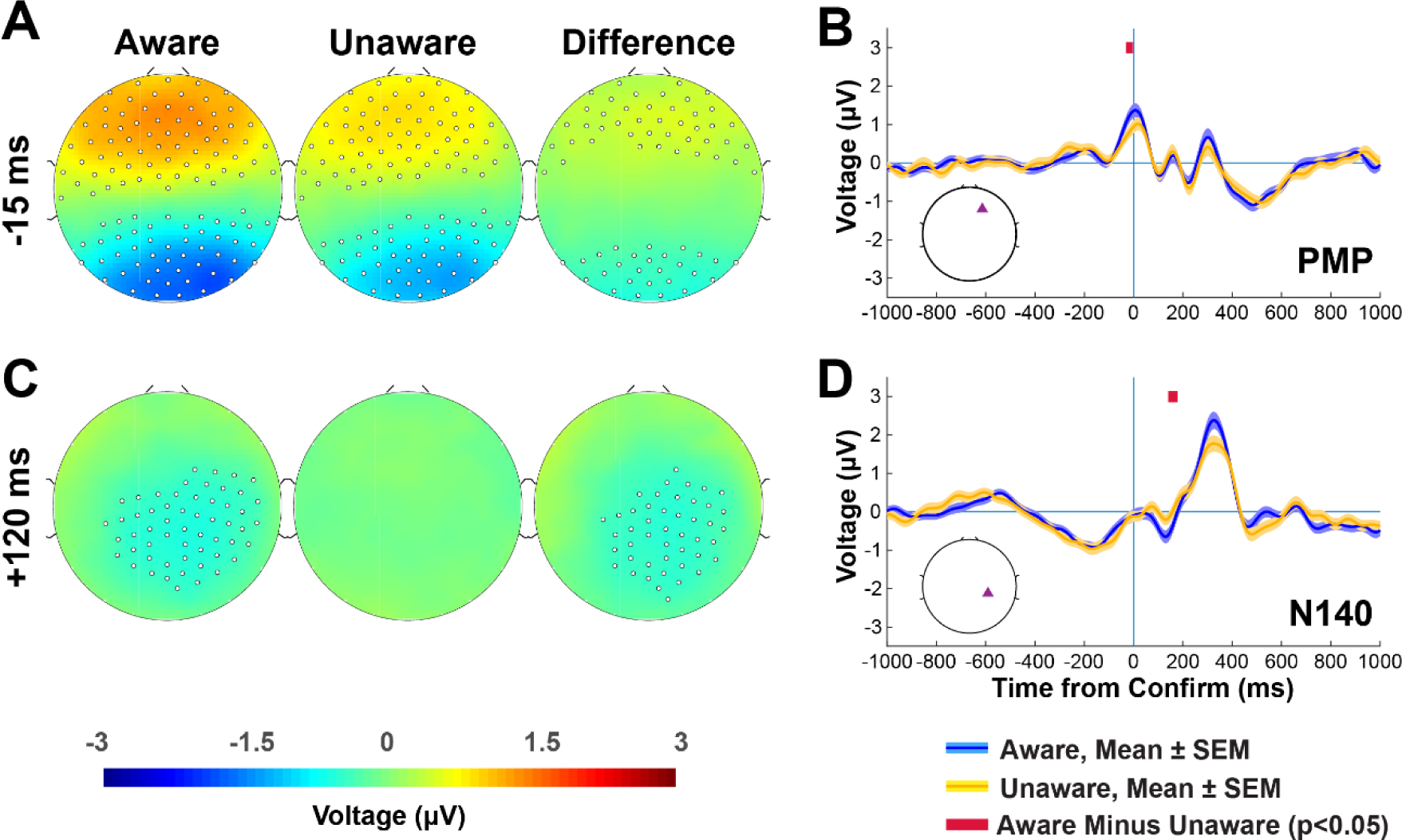
Candidate potentials for AoA. A. Representative topoplot for the pre-movement positivity (PMP). Statistically significant electrodes as determined by spatiotemporal cluster permutation analyses (see Methods – hdEEG spatiotemporal analyses) are shown for aware, unaware, and aware minus unaware conditions. Times indicated are relative to the confirmation of the action. B. Individual timecourse of PMP with SEM. Blue traces indicate aware actions, while yellow traces indicate unaware actions. Crimson timepoints indicate statistically significant timepoints as determined by spatiotemporal criteria (see Methods – hdEEG spatiotemporal analyses). C. Representative topoplot for the somatosensory awareness potential (N140). D. Individual timecourse of N140 with SEM.

### Event Related Potentials – Precursors and Consequences

In addition to the potentials observed in the periphery of action, we also observed volitional precursors occurring in long timescales before (beyond 200 ms pre-action) and attentional consequences in long timescales after (beyond 200 ms post-action) action. Pre-action, we observed a robust negative deflection in scalp voltage over parietal and occipital regions for both aware and unaware actions (**Fig 3C**), beginning ∼600 ms prior to action and peaking ∼150 ms prior to action performance (**Fig 3B**). The magnitude of this potential did not differ between aware and unaware actions. The timing of this potential, the largest observed ERP in magnitude and duration in our experiment, resembles that of the late BP, generated by primary motor cortices[21]. Notably, the parietal and occipital distribution of this potential does not align with the late BP’s spatial distribution over supplementary and primary somatomotor regions. Prior to this potential, we have noted a novel potential, the pre-readiness positivity (PR+), over parietal and occipital regions (**Fig 3A**). This magnitude of the potential was greater in unaware actions compared to aware actions. To ensure that the PR+ was not the result of carry-over from the previous move, we analyzed the mouse click associated with the quizzed move, as well as the confirmation preceding the selection of the quizzed move. We found no differences in these between aware and unaware actions, suggesting that the PR+ was not the result of these action types **(Fig S3)**. In late time periods following action, we observed the well-known postperceptual attention-related potential, the P300[55-59]. As with the N140, the P300 also peaked over parietal and occipital regions, ∼350 ms following the action, and was also greater in amplitude for aware actions (**Fig 3D, E**).

**Figure 3.**
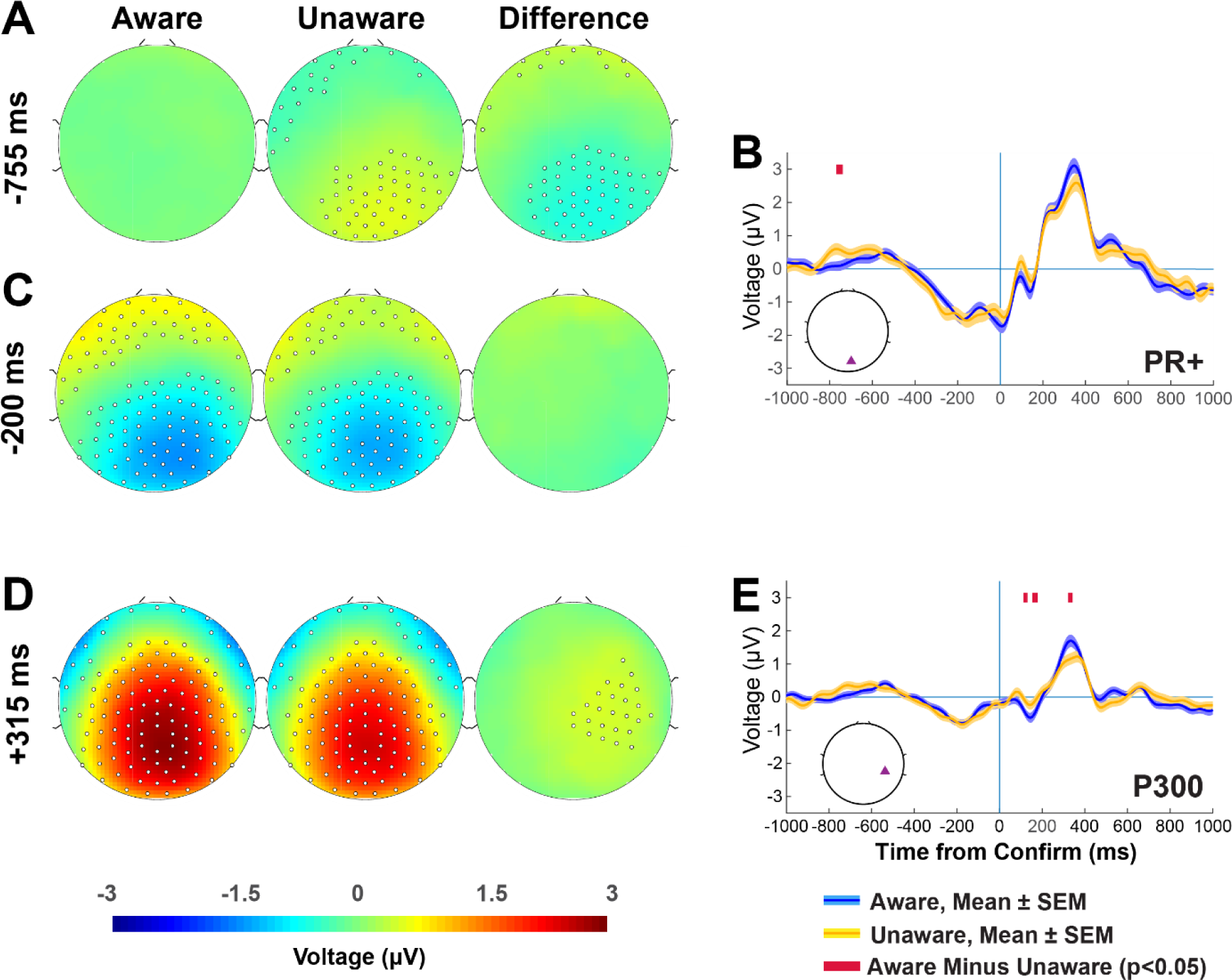
Precursors and consequences of action. A. Representative topoplots for the pre-readiness positivity (PR+). Statistically significant electrodes as determined by spatiotemporal cluster permutation analyses (see Methods – hdEEG spatiotemporal analyses) are shown for aware, unaware, and aware minus unaware conditions. B. Individual timecourse of PR+. Blue traces indicate aware actions, while yellow traces indicate unaware actions. Crimson timepoints indicate statistically significant timepoints as determined by spatiotemporal criteria (see Methods – hdEEG spatiotemporal analyses). C. Representative topoplots of the characteristic pre-action negativity. D. Representative topoplots of the P300. E. Individual timecourse of P300 with SEM.

### Time-Frequency Analyses

To further understand the links between AoA and both volition and perception, we performed time-frequency analyses on our EEG data (**Fig 4**). We found that prior to aware actions, there was a greater event-related desynchronization in the alpha (8-12 Hz) range over frontal and occipital regions (**Fig 4A**). Alpha ERD in aware actions began ∼1000 ms prior to action, peaking at the performance of the action (**Fig 4B**). We also noted event-related desynchronization in beta (12-30 Hz) range over right somatomotor regions preceding aware actions, a finding consistent with previous literature on action generation (**Fig 4C**). Beta ERD in aware actions began ∼150 ms prior to action, and as with alpha ERD, peaked at action performance (**Fig 4D**). Following action, we noted an increase in theta power (4-8 Hz) throughout all electrodes following both aware and unaware actions; however, the magnitude of the increase in theta over midfrontal regions specifically was greater for aware actions compared to unaware actions (**Fig 4E**). The increase in theta power began immediately following action for both aware and unaware actions, and peaked ∼300 ms following aware actions and ∼250 ms following unaware actions (**Fig 4F**). Lastly, we observed a robust post-movement beta rebound throughout all electrodes in both aware and unaware actions (**Fig 4G**), beginning at action and reaching steady state ∼600 ms post-action. The magnitude of the post-movement beta rebound in frontal regions following aware actions exceeded that of unaware actions, from ∼1400 ms post-action onwards (**Fig 4H**).

**Figure 4.**
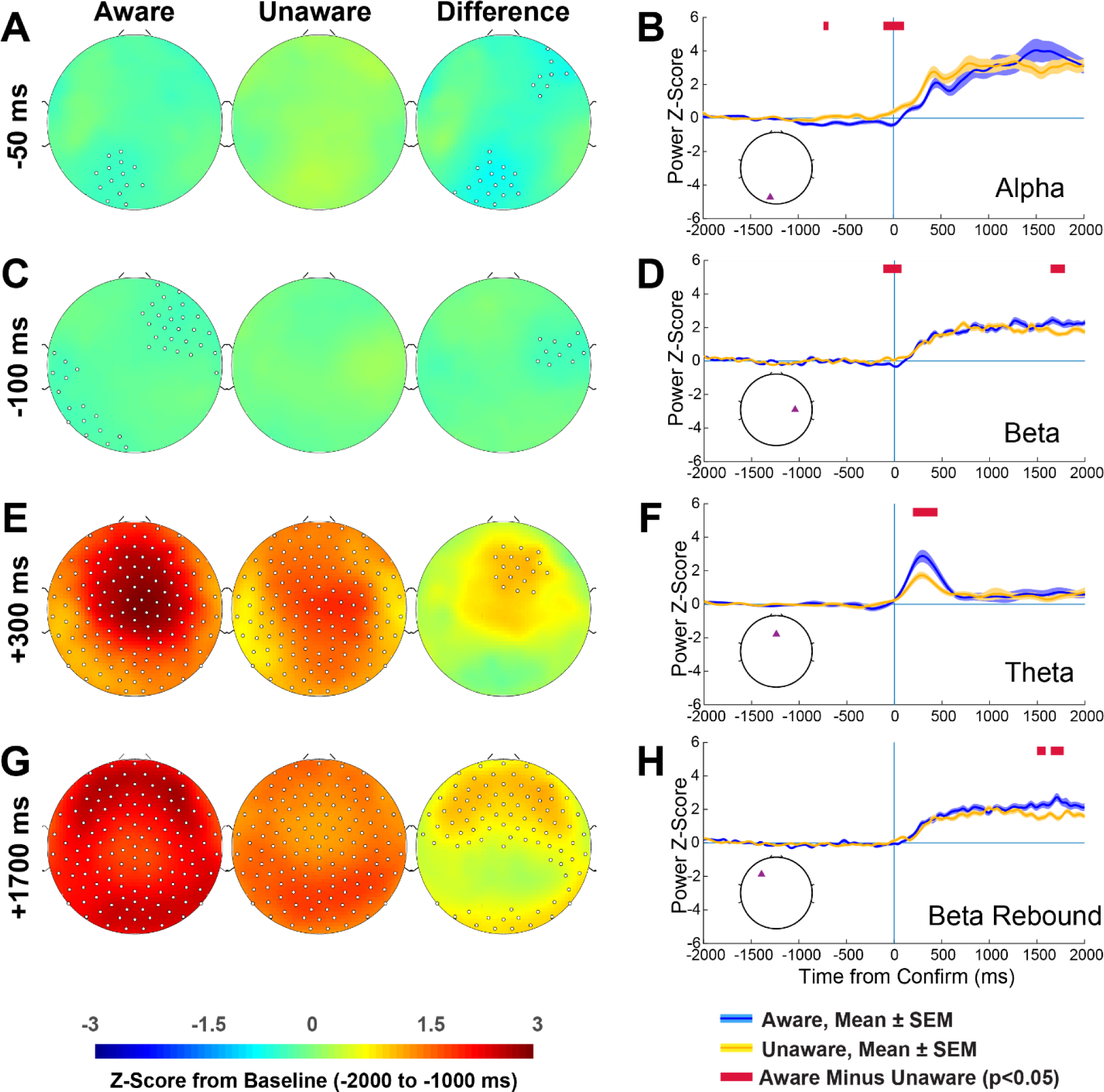
Time frequency analyses. A. Representative topoplots demonstrating pre-action alpha event-related desynchronization (ERD). Statistically significant electrodes as determined by spatiotemporal cluster permutation analyses (see Methods – hdEEG spatiotemporal analyses) are shown for aware, unaware, and aware minus unaware conditions. B. Individual timecourse of alpha ERD shown with SEM. Blue traces indicate aware actions, while yellow traces indicate unaware actions. Crimson timepoints indicate statistically significant timepoints as determined by spatiotemporal criteria (see Methods – hdEEG spatiotemporal analyses). C. Representative topoplots demonstrating pre-action beta ERD. D. Individual timecourse of beta ERD shown with SEM. E. Representative topoplots demonstrating post-action midfrontal theta. F. Individual timecourse with midfrontal theta shown with SEM. G. Representative topoplots showing post-movement beta rebound (PMBR). H. Individual timecourse with PMBR shown with SEM.

### Eye Metrics - Immediate

To assess the relationship between perceptual awareness and AoA, subjects underwent pupillometry and eye tracking while performing the Rush Hour task. Following the confirmation of a move and subsequent disappearance of the board in preparation for a quiz, a consistent increase in blink rate was observed, beginning ∼200 ms post-action and peaking ∼500 ms post-action **(Fig 5A)**. This increase was greater in magnitude for aware actions compared to unaware actions. Post-action increases in pupil diameter (peaking ∼1300 ms post-action) and saccade rates (peaking at ∼350 ms post-action) were observed in both aware and unaware actions, and did not differ in magnitude **(Fig 5B, C)**.

**Figure 5.**
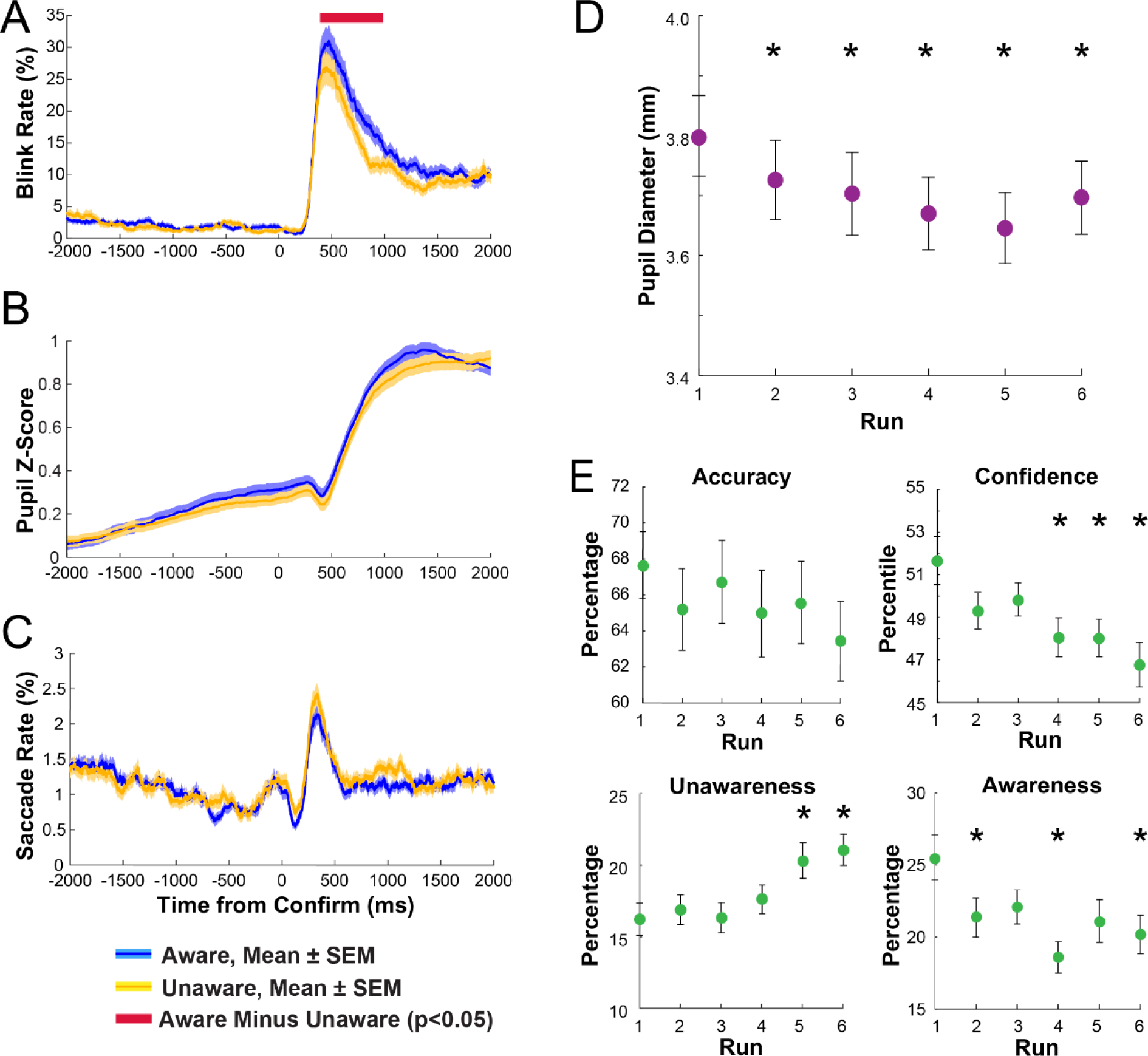
Action-related eye metrics and long-term behavioral and physiological metrics. A. Blink rates following the confirmation of an action. B. Z-scored pupil diameter before and after the confirmation of an action. C. Saccade rates before and after confirmation of an action. Blue traces indicate aware actions, while yellow traces indicate unaware actions. Crimson timepoints indicate statistically significant timepoints as determined by spatiotemporal criteria (see Methods – EyeLink temporal analyses) D. Run-by-run averages of raw pupil diameter. Significance of run-by-run findings was assessed with a one-sample t-test of the current run against Run 1 corrected with the Benjamini-Hochberg procedure. E. Run-by-run averages of quiz accuracy, confidence percentile, awareness rate, and unawareness rate with SEM.

### Behavioral and Physiological Metrics – Long Term

To assess the presence of physiological metrics of fatigue across runs, we analyzed pre-action pupil diameter by run (**Fig. 5D**). We calculated the average pupil diameter one second prior to action (−1000 ms to 0) in all trials, regardless of awareness designation, and found consistent, statistically significant decreases in pupil diameter with each succeeding run. Pupil diameter peaked in Run 1 (mean pupil diameter: 3.8 mm) and fell to its lowest in Run 5 (mean pupil diameter: 3.6 mm).

To assess the effects of fatigue and arousal on the task, we assessed the averages of four key metrics at the whole-run level across the six runs of the experiment: awareness (percentage of aware quizzes), unawareness (percentage of unaware quizzes), quiz accuracy (move identification), and confidence (percentile)(**Fig 5E**). We found that there was a trend towards decreased accuracy across the six runs. As compared to Run 1, we observed statistically significant decreases in awareness in later runs (Runs 2, 4, 6), decreases in confidence (Runs 4, 5, 6), and increases in unawareness rate (Runs 5, 6).

To determine if awareness and unawareness was a sustained state, we looked at whether awareness levels remained the same in successive quizzed moves. The median time period between quizzes was 46.2 seconds, allowing us to observe the maintenance of AoA across multiple minutes. We found that given a quizzed move was aware, the three quizzed moves prior were more likely to also have been aware as compared to if the given move was unaware (with the current quizzed move as move *n*, *n-3*: 22.8% aware if aware vs 16.4% aware if unaware, *n-2*: 22.6% aware if aware vs 14.5% aware if unaware, *n-1* 21.0% aware if aware vs 14.0% aware if unaware). We also found the same for the two quizzed moves following the current move (*n+1*: 20.1% aware if aware vs. 13.3% aware if unaware, *n+2*: 20.6% aware if aware vs. 13.1% aware if unaware. We found that awareness rates for quizzed moves beyond *n-3* and *n+3* generally did not differ regardless of whether move *n* was aware or unaware (with the exception of *n-5*: 22.2% aware if aware vs. 18.3% aware if unaware). Our findings suggest that awareness is a sustained state over tens of seconds (**Fig. S4**).

## Discussion

Our behavioral paradigm has demonstrated that it can reliably separate aware actions from unaware ones, with high rates of behavioral validation. This has enabled us to study the spatiotemporal dynamics of neural activity underlying AoA. We have found clear differences pre- and post-action, in behavioral, pupillometric, ERP, and time-frequency domains between aware and unaware actions. Overall, many ERPs we observed, the PMP, N140, and P300, have all been observed in prior work on perceptual awareness and volitional action. The presence of a heightened PMP prior to action suggests increased pre-action volitional monitoring in AoA, while the presence of a heightened N140 suggests that post-action somatosensory processes allow for the report of action. Our novel pre-action potential, the PR+, suggests that aware and unaware actions may differ in neural activity up to 800 ms pre-action. Our observed negativity 600 ms prior to action showed temporal but not spatial resemblance to the late BP, suggesting it may be a separate action-related process. This was observed regardless of if an action was aware or unaware, suggesting that negative cortical potentials may be common across action-related paradigms. Our time-frequency analyses further indicated that common neural patterns may underlie both volition and perception, and our findings of pre-action event-related desynchronization in the alpha/beta band also suggest that neural mechanisms which are common to both volition and perception are present during aware actions.

The report-based and time between action and quiz raises the question of whether the observed effects are purely the result of memory-related neural mechanisms. It is certain that given sufficient time, such as the course of minutes or hours, the ability to report a move will diminish. However, we observe differences in physiological metrics well before and within the bounds of iconic memory (a few hundred milliseconds)[60, 61]. Furthermore, the time delay between the action performed and the presentation of a quiz remains within the bounds of short-term memory, measured on the scale of tens of seconds[62, 63]. Thus, our findings cannot be purely attributable to memory-related mechanisms.

Our findings also suggest that long timescale, post-action reflection plays a role in maintaining AoA and the ability to report an action. We found that the P300, a later potential, was heightened following aware actions, suggesting that AoA requires post-action attention and preparation to report. In addition to our study, the combination of the N140 and P300 is observed across multiple somatosensory task paradigms, suggesting that the P300 is an attentional consequence of somatosensation[64-66]. We also found increases in midfrontal theta in aware actions, a reflective mechanism seen in cognitive control tasks involving a motor component[67, 68]. Our findings also provide support to other studies which have highlighted the role of midfrontal theta in successful navigation[69-71]. In the longer post-action timescales (>1500 s), we noted increases in post-movement beta rebound following aware actions. Post-movement beta rebound is generated by inhibition of motor cortex and preparation for sensory feedback of action, a long-term consequence of action[72-75].

The eye metrics seen in our finding indicate that spontaneous blink rate may be the most indicative of AoA. Previous primate research has suggested that somatosensory areas, in conjunction with visual areas, are important contributors to spontaneous eyeblinks, indicating possible overlap with AoA-related brain regions[76]. Additionally, blink rates are known to correlate with dopamine levels in the brain, suggesting a dopaminergic mechanism for AoA[77]. Saccades, on the other hand, are hallmarks of visual attention, with direct links from primary visual cortex[78, 79]. The fact that we found no difference in saccade rates between aware and unaware actions suggests that the visual modality is less important than somatosensation in maintaining AoA.

We found behavioral patterns and physiological measurements suggesting a heavy role of arousal and fatigue in the maintenance of AoA. These patterns and biological signals occurred over the course of multiple seconds or even over the course of minutes. This sampling frequency allowed us to look at state-based factors of awareness within a typical attention span. Our sequential quiz-to-quiz analyses suggest that periods of awareness are maintained for ∼4.5 minutes, nearly half of an entire run. Our run-by-run behavioral findings of decreased confidence, accuracy, and awareness, as well as increased unawareness rates, suggest that long-term decreases in arousal compromise AoA. Our run-by-run decreases in pupil diameter support our behavioral findings, with physiological evidence for decreased arousal across runs.

While our study has encapsulated much of the action patterns and planning seen in naturalistic AoA, it still remains to be seen if the kinesthetics underlying an action affect AoA. Our study has probed navigation, only one naturalistic situation under which AoA is lost. Other effortful actions, such as those involving physical fatigue, or grasping while obstructed may show diverging neural pathways[80-82]. Furthermore, a whole body experience may differ from a simple point-and-click game, especially when coordinating multiple limbs[83, 84]. Additionally, our study has also probed only the cortical mechanisms of AoA. Given that action is known to be generated in the cerebellum, brainstem, basal ganglia, and other subcortical regions, further studies with fMRI and other imaging modalities are needed[85-88]. Lastly, our behavioral experiment is readily adaptable to interventional modalities which have been tested on motor regions, such as transcranial magnetic stimulation and transcranial alternating current stimulation[89-92]. These studies will allow causal neuroanatomical determinants of AoA to be conclusively identified.

The overlap between findings in our clinically normal population and populations with impaired AoA is notable. Two of our time-frequency findings, midfrontal theta and beta desynchronization, are known to be diminished in patients with Parkinson’s disease[93-95]. Furthermore, pre-movement event-related desynchronization and post-movement beta rebound are attenuated in amylotrophic lateral sclerosis, and beta hypersynchronization is noted in dystonia[94, 96]. This suggests that the aberrant AoA seen in these movement disorders may be related to loss of control over normal awareness processes. Further testing of our novel behavioral paradigm in clinical populations will be able to validate this hypothesis, and determine its value as a part of diagnostic batteries.

Our findings have given a definitive answer to the Two Williams Debate. Given our findings of both pre- and post-action differences in neural activity between aware and unaware actions, we conclude that it is impossible to ascribe the neural bases of AoA to purely *a priori*, volitional or purely *a posteriori*, perceptual mechanisms. Thus, both perspectives held in the Two Williams Debate hold merit in explaining the mechanisms underlying action. Furthermore, our longer-term findings in the precursors and consequences of action suggest that the dichotomous view of action mechanisms as either pre- or post-action is incomplete. These long-term findings demonstrate that beyond its characterization with dynamic, transient neural activity, AoA is also characterized by static, long-term states. Further investigation of AoA through the perspective of individual events and overarching states in tandem will help reveal the richness of neural mechanisms underlying voluntary action.

## Acknowledgements

We thank Dr. Linda Mayes, Dr. Helena Rutherford, Dr. Todd Constable, the Yale Child Study Center, and the Yale Magnetic Resonance Research Center for their assistance in the process of EEG data acquisition in their respective facilities. We also thank Dr. Ayushe Sharma for providing immense assistance in the manuscript and figure preparation process.

---

**Supplementary Figure 1.**
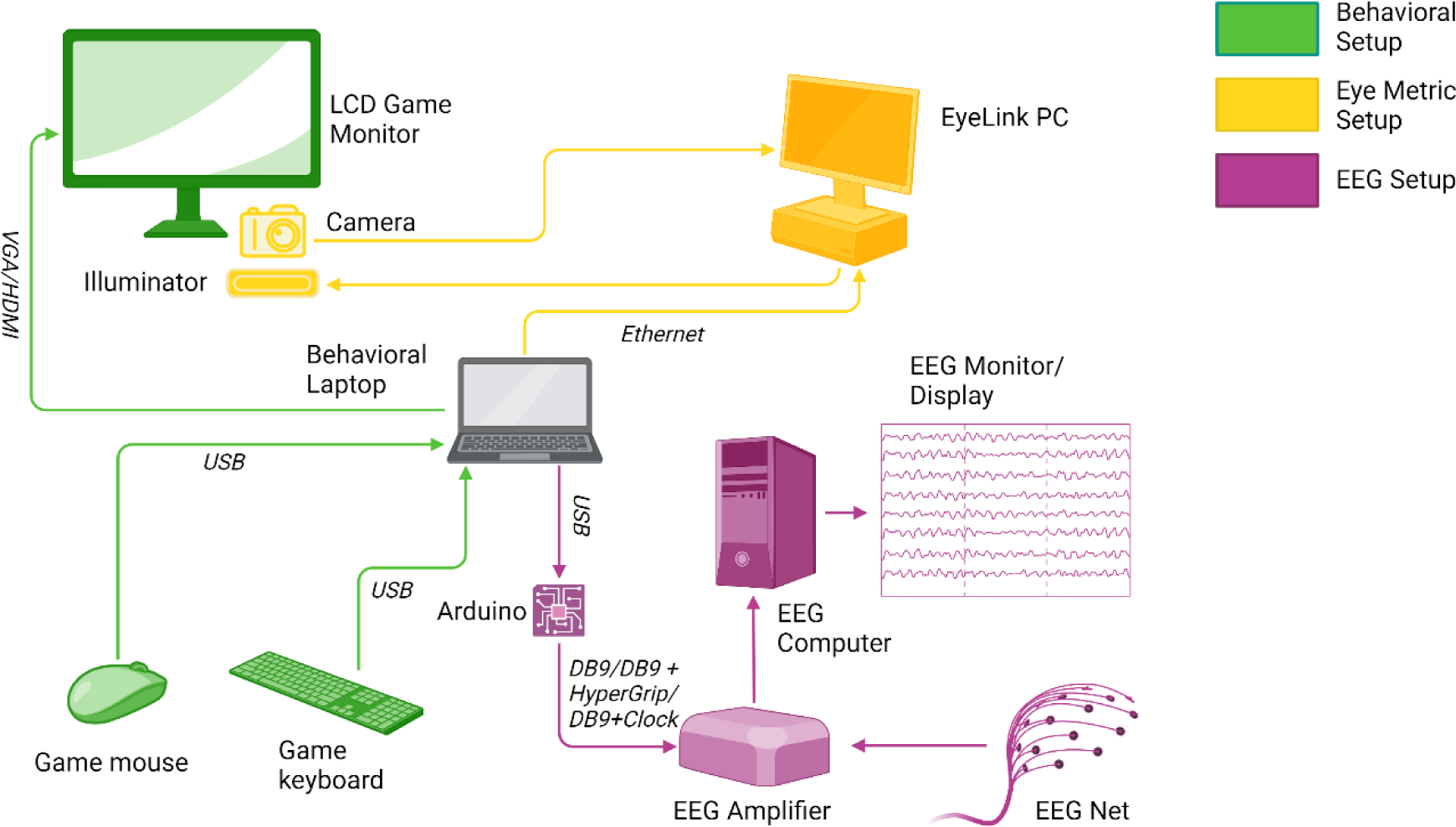
Experimental setup. Subjects performed the task using items shown in green. Pupillometry-related items are shown in yellow, and EEG-related items are shown in purple. The Arduino to amplifier connection was established with three methods (1) direct DB9 connection (N = 30 subjects), (2) DB9 to clock box (N = 7 subjects), and (3) DB9 to hypergrip (N = 20 subjects).

**Supplementary Figure 2.**
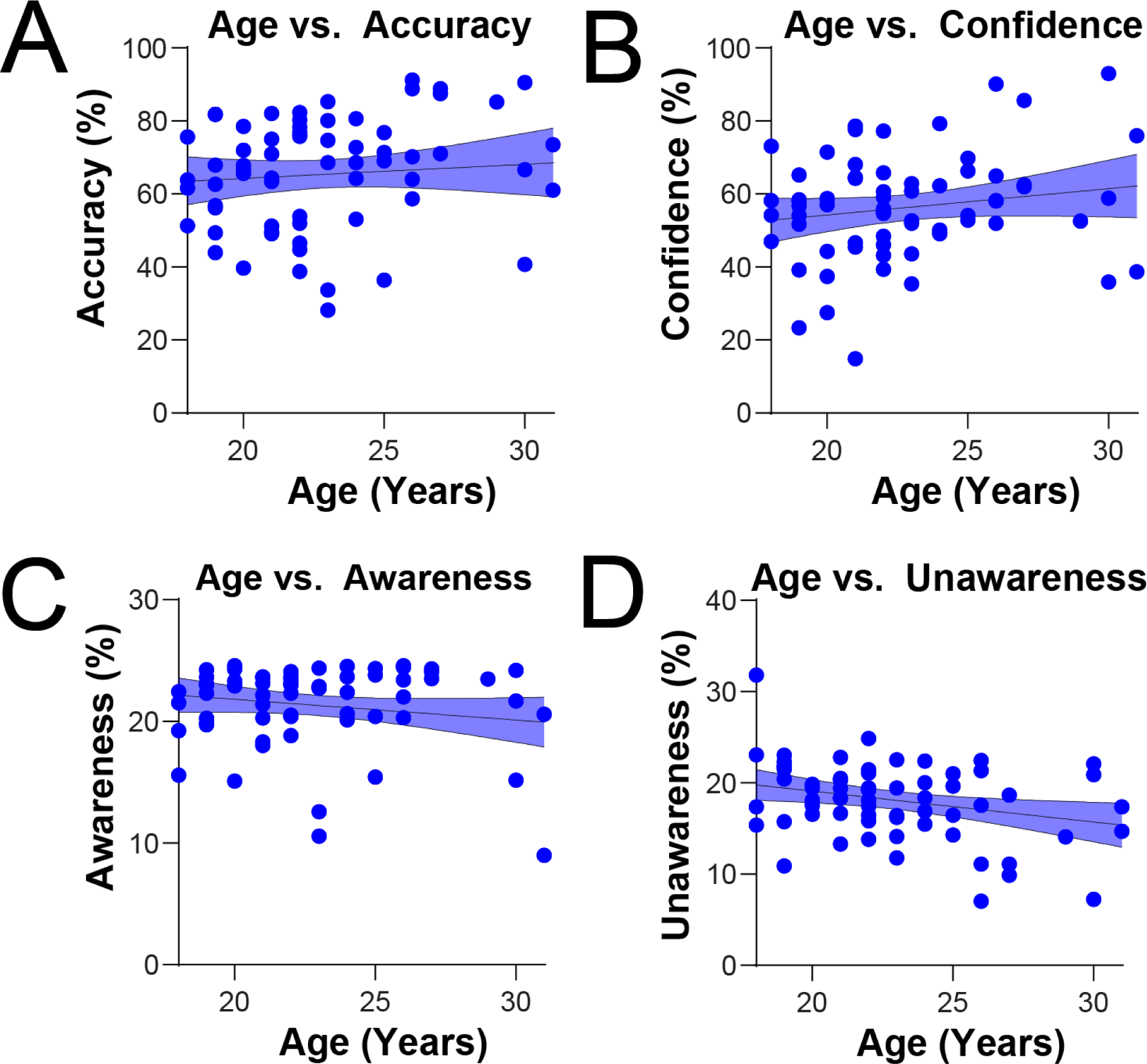
Relationship between age and key metrics of A. Accuracy (r = 0.09 p = 0.47), B. Confidence (r = 0.18, p = 0.15), C. Awareness (r = −0.18, p = 0.15) and D. Unawareness (r = −0.29*, p = 0.02*).

**Supplementary Figure 3.**
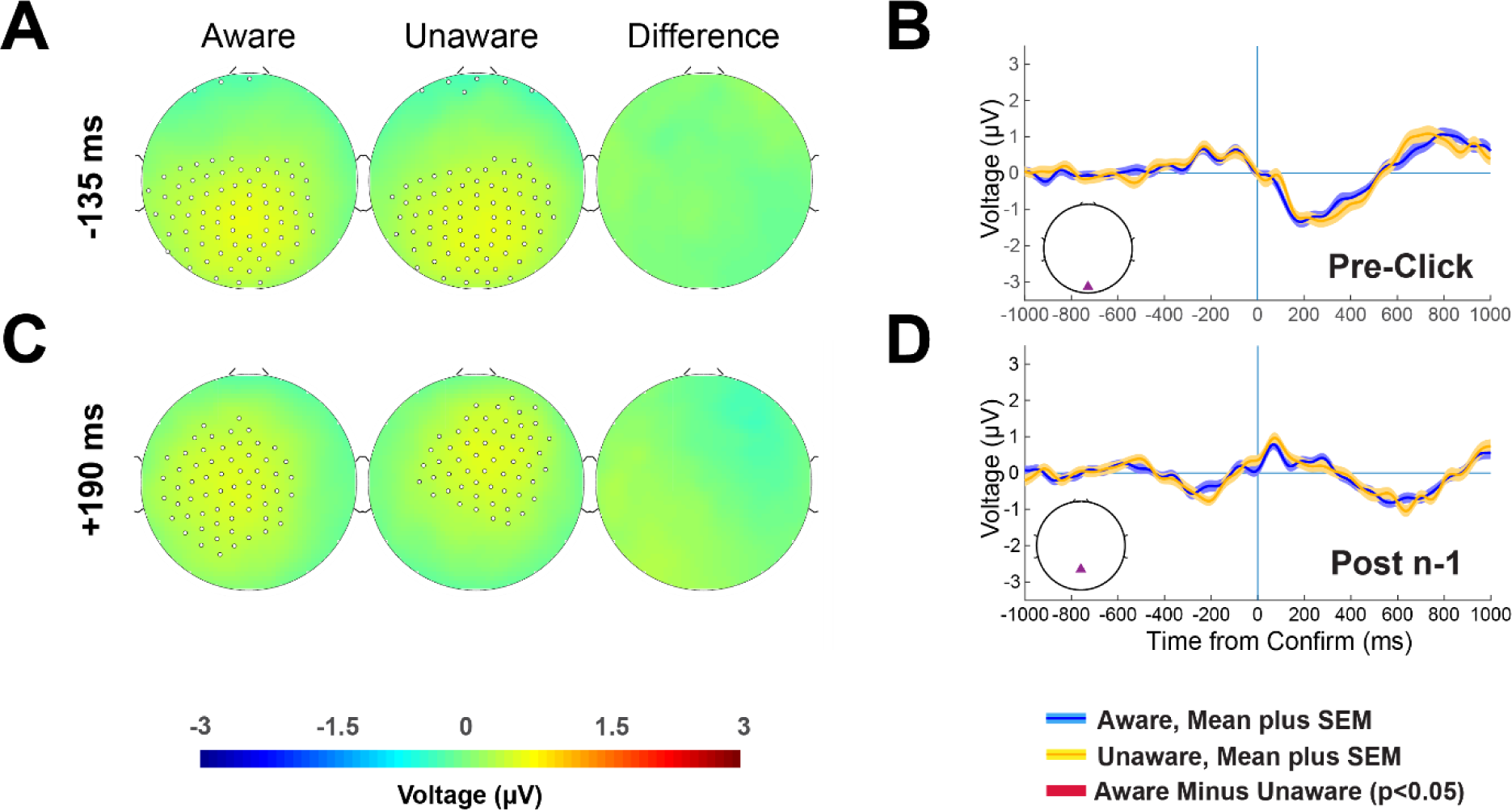
Assessments of other epochs for potential leakage to the PR+. A. Representative topoplots for the move selection associated with the quizzed action. B. Individual timecourse of the move selection epoch with SEM. C. Representative topoplots for the confirmation of the move immediately preceding a quizzed action. D. Individual timecourse of the confirmation of the move preceding a quizzed action with SEM. Blue traces indicate aware actions, while yellow traces indicate unaware actions. Crimson timepoints indicate statistically significant timepoints as determined by spatiotemporal criteria (see Methods – hdEEG spatiotemporal analyses).

**Supplementary Figure 4.**
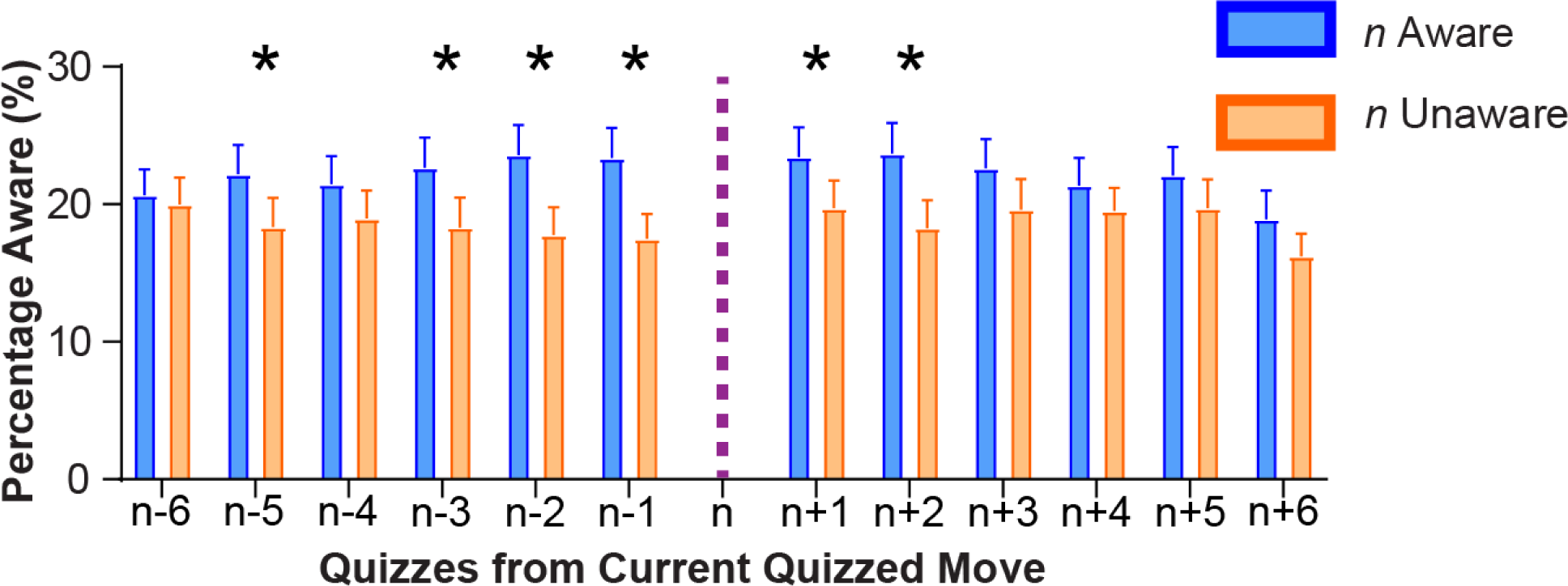
Awareness rates surrounding a move, designated move *n*, which is either aware (blue) or unaware (orange). Statistical significance was assessed with a one sample paired t-test with Benjamini-Hochberg FDR correction.

**Supplementary Table S1.** Awareness and unawareness by block color. Block identity association with awareness levels. Red indicates the target block which must be moved to solve the puzzle, while white indicates any obstructing block. Unvalidated answers were considered those that were either correct and low confidence, or incorrect and high confidence. χ^2^ tests compared awareness and unawareness for the target red block versus other white blocks. Three categorizations were used: 1. Two category (aware vs. unaware) 1. Three category (aware, unaware, other) and 2. Five-category (aware, unaware, mid-high confidence, mid-low confidence, unvalidated). The “other” category was defined as any quiz in either mid-high, mid-low, or unvalidated categories.

|  | *Red Blocks<sup>1</sup>* | *White Blocks<sup>1</sup>* |
| --- | --- | --- |
|  | N (%) | N (%) |
| Aware<sup>2</sup> | 163 (21.0%) | 1,614 (21.2%) |
| Unaware<sup>2</sup> | 156 (20.0%) | 1,354 (17.8%) |
| Mid-High<sup>2</sup> | 201 (25.8%) | 1,870 (24.5%) |
| Mid-Low<sup>2</sup> | 187 (24.0%) | 1,977 (25.9%) |
| Unvalidated<sup>2</sup> | 72 (9.2%) | 812 (10.6%) |
| Total | 779 (100%) | 7,627 (100%) |

| | $\chi^2$ | *p-value* |
| --- | --- | --- |
| Aware vs Unaware | 1.25 | 0.26 |
| Aware vs Unaware vs<br>(Mid-High, Mid-Low<br>or Unvalidated) | 2.53 | 0.28 |
| Aware vs Unaware vs<br>Mid-High vs Mid-Low vs Unvalidated | 4.86 | 0.30 |

<sup>1</sup>Total number of Red or White blocks quizzed across all participants, and percent of total.
<sup>2</sup>Definitions as follows, based on Figure 1C, D: Aware trials have correct move identification on multiple choice and confidence percentile >75%; Unaware trials have incorrect move identification on multiple choice and confidence percentile <25%; Mid-High trials have confidence percentile >50% and ≤75% (regardless of answer on multiple choice); Mid-Low trials have confidence percentile ≥25% and ≤50% (regardless of answer on multiple choice); Unvalidated trials have either incorrect move identification on multiple choice and confidence percentile >75% or correct move identification on multiple choice and confidence percentile <25%.

**Supplementary Table S2.**
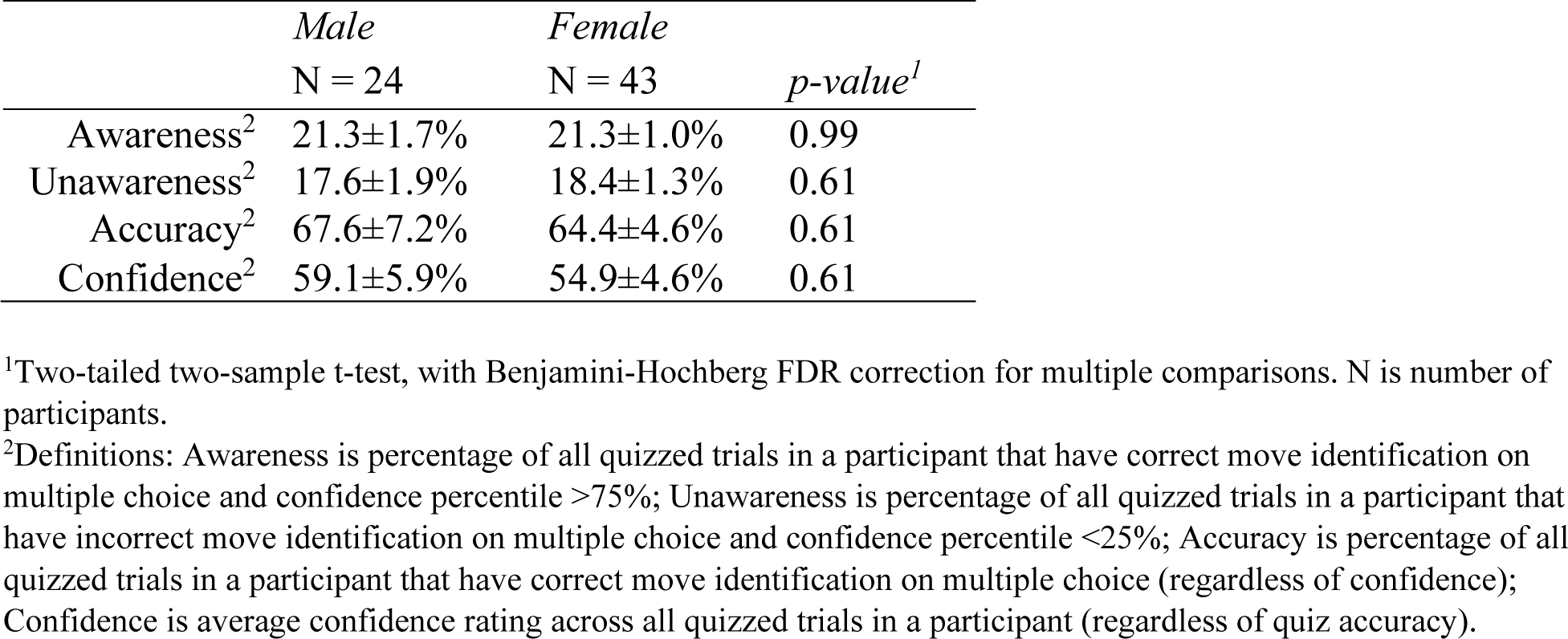
Relationship between key behavioral metrics and sex. Relationship between sex and four key metrics of awareness, unawareness, quiz accuracy, and quiz confidence. Sex was tested with a two-tailed two-sample t-test (p<0.05).

**Supplementary Table S3.** Relationship between key behavioral metrics and testing day. Relationship between testing day and four key metrics of awareness, unawareness, quiz accuracy, and quiz confidence. day was tested with a two-tailed paired t-test (p<0.05).

|  | *Day 1*<br>*N=61* | *Day 2*<br>*N=61* | *p-value*<sup>1</sup> |
| --- | --- | --- | --- |
| Awareness<sup>2</sup> | 21.7±1.1% | 22.3±1.1% | 0.50 |
| Unawareness<sup>2</sup> | 17.4±1.0% | 17.0±1.2% | 0.61 |
| Accuracy<sup>2</sup> | 64.9±4.1% | 67.6±4.5% | 0.15 |
| Confidence<sup>2</sup> | 57.0±3.7% | 57.9±4.0% | 0.98 |

<sup>1</sup>Two-tailed paired t-test, with Benjamini-Hochberg FDR correction for multiple comparisons. N is number of participants.
<sup>2</sup>Definitions: Awareness is percentage of all quizzed trials in a participant that have correct move identification on multiple choice and confidence percentile >75%; Unawareness is percentage of all quizzed trials in a participant that have incorrect move identification on multiple choice and confidence percentile <25%; Accuracy is percentage of all quizzed trials in a participant that have correct move identification on multiple choice (regardless of confidence); Confidence is average confidence rating across all quizzed trials in a participant (regardless of quiz accuracy).

**Supplementary Table S4.**
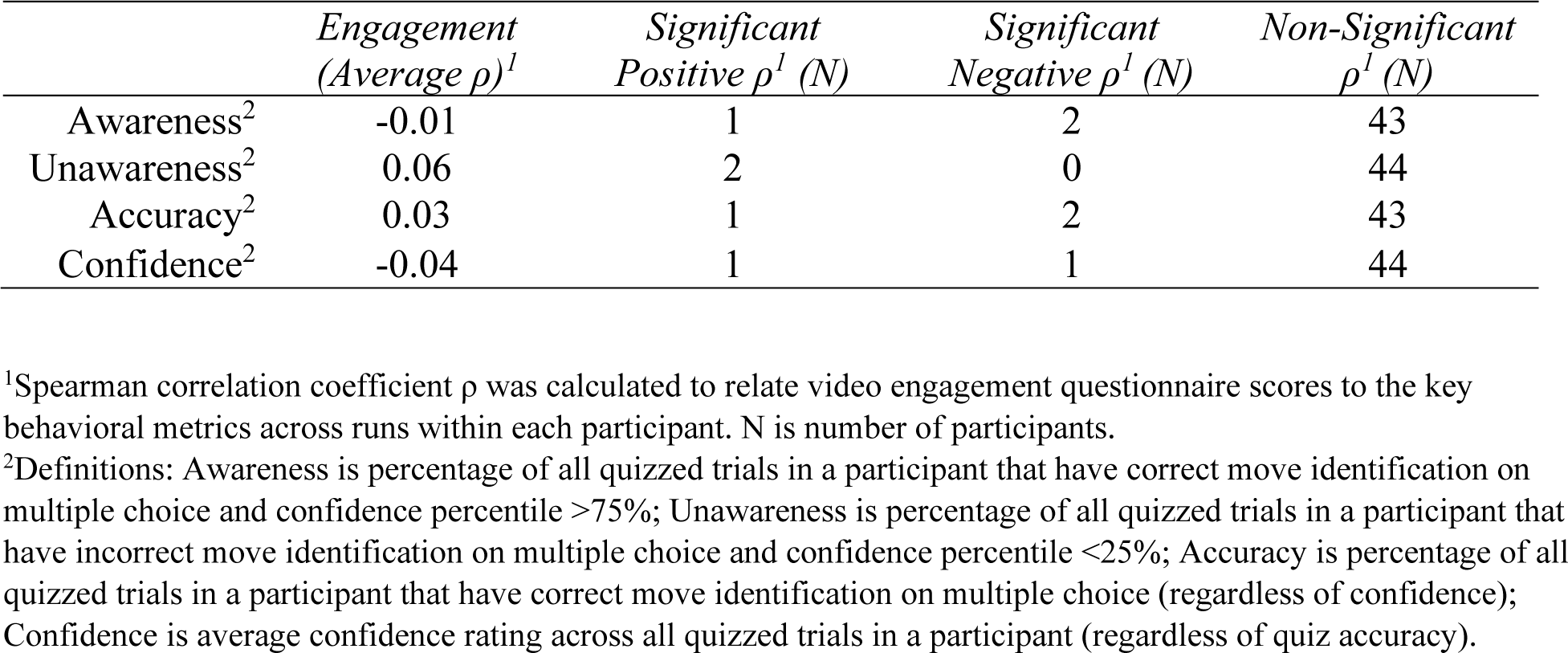
Relationship between key behavioral metrics and video engagement. Relationship between video engagement rating and within-run averages of four key metrics of awareness, unawareness, quiz accuracy, and quiz confidence (N = 46). Engagement correlations within subject were assessed with a Spearman correlation coefficient. The number of statistically significant and non-significant within-subject correlations is listed.

|  | *Engagement*<br>*(Average $\rho$)<sup>1</sup>* | *Significant*<br>*Positive $\rho^1$ (N)* | *Significant*<br>*Negative $\rho^1$ (N)* | *Non-Significant*<br>*$\rho^1$ (N)* |
| --- | --- | --- | --- | --- |
| Awareness<sup>2</sup> | -0.01 | 1 | 2 | 43 |
| Unawareness<sup>2</sup> | 0.06 | 2 | 0 | 44 |
| Accuracy<sup>2</sup> | 0.03 | 1 | 2 | 43 |
| Confidence<sup>2</sup> | -0.04 | 1 | 1 | 44 |

<sup>1</sup>Spearman correlation coefficient $\rho$ was calculated to relate video engagement questionnaire scores to the key behavioral metrics across runs within each participant. N is number of participants.
<sup>2</sup>Definitions: Awareness is percentage of all quizzed trials in a participant that have correct move identification on multiple choice and confidence percentile >75%; Unawareness is percentage of all quizzed trials in a participant that have incorrect move identification on multiple choice and confidence percentile <25%; Accuracy is percentage of all quizzed trials in a participant that have correct move identification on multiple choice (regardless of confidence); Confidence is average confidence rating across all quizzed trials in a participant (regardless of quiz accuracy).

**Supplementary Table S5.** Relationship between key behavioral metrics and video familiarity. Relationship between video familiarity rating and within-run averages of four key metrics of awareness, unawareness, quiz accuracy, and quiz confidence (N = 45). Engagement correlations within subject were assessed with a Spearman correlation coefficient. The number of statistically significant and non-significant within-subject correlations is listed.

|  | *Familiarity*<br>*(Average $\rho$)<sup>1</sup>* | *Significant*<br>*Positive ρ<sup>1</sup>* | *Significant*<br>*Negative ρ<sup>1</sup>* | *Non-*<br>*Significant ρ<sup>1</sup>* |
| --- | --- | --- | --- | --- |
| Awareness<sup>2</sup> | 0.02 | 2 | 2 | 41 |
| Unawareness<sup>2</sup> | 0.08 | 2 | 0 | 43 |
| Accuracy<sup>2</sup> | 0.00 | 1 | 1 | 43 |
| Confidence<sup>2</sup> | 0.00 | 1 | 2 | 42 |

<sup>1</sup>Spearman correlation coefficient $\rho$ was calculated to relate video familiarity questionnaire scores to the key behavioral metrics across runs within each participant. N is number of participants.
<sup>2</sup>Definitions: Awareness is percentage of all quizzed trials in a participant that have correct move identification on multiple choice and confidence percentile >75%; Unawareness is percentage of all quizzed trials in a participant that have incorrect move identification on multiple choice and confidence percentile <25%; Accuracy is percentage of all quizzed trials in a participant that have correct move identification on multiple choice (regardless of confidence); Confidence is average confidence rating across all quizzed trials in a participant (regardless of quiz accuracy).

## References

1. Cerezuela, G.P., et al., Wertheim’s hypothesis on ‘highway hypnosis’: empirical evidence from a study on motorway and conventional road driving. Accid Anal Prev, 2004. 36(6): p. 1045–54.

2. Wertheim, A.H., Explaining highway hypnosis: Experimental evidence for the role of eye movements. Accident Analysis & Prevention, 1978. 10(2): p. 111–129.

3. Frith, C.D., S. Blakemore, and D.M. Wolpert, Explaining the symptoms of schizophrenia: abnormalities in the awareness of action. Brain Res Brain Res Rev, 2000. 31(2-3): p. 357–63.

4. Tinaz, S., et al., Insula as the Interface Between Body Awareness and Movement: A Neurofeedback-Guided Kinesthetic Motor Imagery Study in Parkinson’s Disease. Front Hum Neurosci, 2018. 12: p. 496.

5. Berryessa, C.M., F. Coppola, and G. Salvato, *The potential effect of neurobiological evidence on the adjudication of criminal responsibility of psychopathic defendants in involuntary manslaughter cases.* Psychology, Crime & Law, 2021. 27(2): p. 140–158.

6. Thomaidou, M.A. and C.M. Berryessa, A jury of scientists: Formal education in biobehavioral sciences reduces the odds of punitive criminal sentencing. Behavioral Sciences & the Law, 2022. 40(6): p. 787–817.

7. Hafner, M., Judging homicide defendants by their brains: an empirical study on the use of neuroscience in homicide trials in Slovenia. Journal of Law and the Biosciences, 2019. 6(1): p. 226–254.

8. Jeannerod, M., The origin of voluntary action: history of a physiological concept. C R Biol, 2006. 329(5-6): p. 354–62.

9. Maoz, U., et al., Neural precursors of decisions that matter-an ERP study of deliberate and arbitrary choice. Elife, 2019. 8.

10. Libet, B., et al., Time of conscious intention to act in relation to onset of cerebral activity (readiness-potential). The unconscious initiation of a freely voluntary act. Brain, 1983. 106(Pt 3): p. 623–42.

11. Nann, M., et al., To jump or not to jump - The Bereitschaftspotential required to jump into 192-meter abyss. Sci Rep, 2019. 9(1): p. 2243.

12. Gusso, M.M., et al., More than a feeling: Scalp EEG and eye signals in conscious tactile perception. Conscious Cogn, 2022. 105: p. 103411.

13. Herman, W.X., et al., A Switch and Wave of Neuronal Activity in the Cerebral Cortex During the First Second of Conscious Perception. Cereb Cortex, 2019. 29(2): p. 461–474.

14. Kronemer, S.I., et al., Human visual consciousness involves large scale cortical and subcortical networks independent of task report and eye movement activity. Nat Commun, 2022. 13(1): p. 7342.

15. Christison-Lagay, K.L., et al., The neural activity of auditory conscious perception. BioRxiv, 2023. https://www.biorxiv.org/content/10.1101/2023.01.12.523829v1.full.pdf.

16. Aru, J., et al., Distilling the neural correlates of consciousness. Neurosci Biobehav Rev, 2012. 36(2): p. 737–46.

17. Kim, C.Y. and R. Blake, Psychophysical magic: rendering the visible ‘invisible’. Trends Cogn Sci, 2005. 9(8): p. 381–8.

18. Pares-Pujolras, E., et al., Latent awareness: Early conscious access to motor preparation processes is linked to the readiness potential. Neuroimage, 2019. 202: p. 116140.

19. Schultze-Kraft, M., et al., Preparation and execution of voluntary action both contribute to awareness of intention. Proc Biol Sci, 2020. 287(1923): p. 20192928.

20. Kornhuber, H.H. and L. Deecke, [Changes in the Brain Potential in Voluntary Movements and Passive Movements in Man: Readiness Potential and Reafferent Potentials]. Pflugers Arch Gesamte Physiol Menschen Tiere, 1965. 284: p. 1–17.

21. Schurger, A., et al., What Is the Readiness Potential? Trends Cogn Sci, 2021. 25(7): p. 558–570.

22. Haggard, P., Human volition: towards a neuroscience of will. Nat Rev Neurosci, 2008. 9(12): p. 934–46.

23. Haggard, P. and M. Eimer, On the relation between brain potentials and the awareness of voluntary movements. Exp Brain Res, 1999. 126(1): p. 128–33.

24. Cunnington, R., et al., The preparation and readiness for voluntary movement: a high-field event-related fMRI study of the Bereitschafts-BOLD response. Neuroimage, 2003. 20(1): p. 404–12.

25. Sakata, H., et al., Slow Accumulations of Neural Activities in Multiple Cortical Regions Precede Self-Initiation of Movement: An Event-Related fMRI Study. eNeuro, 2017. 4(5).

26. Bortoletto, M., et al., Pre-motion positivity during self-paced movements of finger and mouth. Neuroreport, 2006. 17(9): p. 883–6.

27. Castro, A., Diaz, F., and van Boxtel, G. J. M., What happens to the readiness potential when the movement is not executed? Neuroreport, 2005. 16(15): p. 1609–1613.

28. Rauchbauer, B., D.M. Pfabigan, and C. Lamm, Event-related potentials of automatic imitation are modulated by ethnicity during stimulus processing, but not during motor execution. Sci Rep, 2018. 8(1): p. 12760.

29. Stephan, K.M., et al., Functional anatomy of the mental representation of upper extremity movements in healthy subjects. J Neurophysiol, 1995. 73(1): p. 373–86.

30. Ball, T., et al., The role of higher-order motor areas in voluntary movement as revealed by high-resolution EEG and fMRI. Neuroimage, 1999. 10(6): p. 682–94.

31. Bode, S., et al., Tracking the unconscious generation of free decisions using ultra-high field fMRI. PLoS One, 2011. 6(6): p. e21612.

32. Dembski, C., C. Koch, and M. Pitts, Perceptual awareness negativity: a physiological correlate of sensory consciousness. Trends Cogn Sci, 2021. 25(8): p. 660–670.

33. Kida, T., et al., Differential modulation of temporal and frontal components of the somatosensory N140 and the effect of interstimulus interval in a selective attention task. Brain Res Cogn Brain Res, 2004. 19(1): p. 33–9.

34. Pavlidou, A., A. Schnitzler, and J. Lange, Distinct spatio-temporal profiles of beta-oscillations within visual and sensorimotor areas during action recognition as revealed by MEG. Cortex, 2014. 54: p. 106–16.

35. Barone, J. and H.E. Rossiter, Understanding the Role of Sensorimotor Beta Oscillations. Front Syst Neurosci, 2021. 15: p. 655886.

36. Espenhahn, S., et al., Movement-related beta oscillations show high intra-individual reliability. Neuroimage, 2017. 147: p. 175–185.

37. Zaretskaya, N. and A. Bartels, Gestalt perception is associated with reduced parietal beta oscillations. Neuroimage, 2015. 112: p. 61–69.

38. Pfurtscheller, G., C. Neuper, and W. Mohl, Event-related desynchronization (ERD) during visual processing. Int J Psychophysiol, 1994. 16(2-3): p. 147–53.

39. Pfurtscheller, G., A. Stancak, Jr., and C. Neuper, Event-related synchronization (ERS) in the alpha band--an electrophysiological correlate of cortical idling: a review. Int J Psychophysiol, 1996. 24(1-2): p. 39–46.

40. Woodman, G.F., et al., Alpha suppression indexes a spotlight of visual-spatial attention that can shine on both perceptual and memory representations. Psychon Bull Rev, 2022. 29(3): p. 681–698.

41. Babiloni, C., et al., Human movement-related potentials vs desynchronization of EEG alpha rhythm: a high-resolution EEG study. Neuroimage, 1999. 10(6): p. 658–65.

42. Fujioka, T. and B. Ross, Auditory processing indexed by stimulus-induced alpha desynchronization in children. Int J Psychophysiol, 2008. 68(2): p. 130–40.

43. Yakovlev, L., N. Syrov, and A. Kaplan, Investigating the influence of functional electrical stimulation on motor imagery related mu-rhythm suppression. Front Neurosci, 2023. 17: p. 1202951.

44. Crone, N.E., A. Sinai, and A. Korzeniewska, High-frequency gamma oscillations and human brain mapping with electrocorticography. Prog Brain Res, 2006. 159: p. 275–95.

45. Hayat, H., et al., Reduced neural feedback signaling despite robust neuron and gamma auditory responses during human sleep. Nat Neurosci, 2022. 25(7): p. 935–943.

46. Neuper, C. and G. Pfurtscheller, Event-related dynamics of cortical rhythms: frequency-specific features and functional correlates. Int J Psychophysiol, 2001. 43(1): p. 41–58.

47. Esterman, M., et al., In the zone or zoning out? Tracking behavioral and neural fluctuations during sustained attention. Cereb Cortex, 2013. 23(11): p. 2712–23.

48. Esterman, M., M.D. Rosenberg, and S.K. Noonan, Intrinsic fluctuations in sustained attention and distractor processing. J Neurosci, 2014. 34(5): p. 1724–30.

49. Simon, A.J., et al., Quantifying attention span across the lifespan. Front Cognit, 2023. 2.

50. Schneider, M., et al., Spontaneous pupil dilations during the resting state are associated with activation of the salience network. Neuroimage, 2016. 139: p. 189–201.

51. DiNuzzo, M., et al., Brain Networks Underlying Eye’s Pupil Dynamics. Front Neurosci, 2019. 13: p. 965.

52. Maki-Marttunen, V., Pupil-based States of Brain Integration across Cognitive States. Neuroscience, 2021. 471: p. 61–71.

53. Kumar, N., Alam, K., and Siddiqi, A. H., Wavelet Transform for Classification of EEG Signal using SVM and ANN. Biomedical & Pharmacology Journal, 2017. 10(4): p. 2061–2019.

54. Groppe, D.M., T.P. Urbach, and M. Kutas, Mass univariate analysis of event-related brain potentials/fields I: a critical tutorial review. Psychophysiology, 2011. 48(12): p. 1711–25.

55. Picton, T.W., The P300 wave of the human event-related potential. J Clin Neurophysiol, 1992. 9(4): p. 456–79.

56. Rohaut, B. and L. Naccache, Disentangling conscious from unconscious cognitive processing with event-related EEG potentials. Rev Neurol (Paris), 2017. 173(7-8): p. 521–528.

57. Rutiku, R., et al., Does the P300 reflect conscious perception or its consequences? Neuroscience, 2015. 298: p. 180–9.

58. Schroder, P., T. Nierhaus, and F. Blankenburg, Dissociating Perceptual Awareness and Postperceptual Processing: The P300 Is Not a Reliable Marker of Somatosensory Target Detection. J Neurosci, 2021. 41(21): p. 4686–4696.

59. Sutton, S., et al., Evoked-potential correlates of stimulus uncertainty. Science, 1965. 150(3700): p. 1187-8.

60. Sperling, G., The information available in brief visual presentations. Psychological Monographs: General and Applied, 1960. 74(11): p. 1–29.

61. Block, N., Perceptual consciousness overflows cognitive access. Trends in Cognitive Sciences, 2011. 15(12): p. 567–575.

62. Shiffrin, R.M. and J.R. Cook, Short-term forgetting of item and order information. Journal of Verbal Learning & Verbal Behavior, 1978. 17(2): p. 189–218.

63. Reitman, J.S., Mechanisms of forgetting in short-term memory. Cognitive Psychology, 1971. 2(2): p. 185–195.

64. Lindenbaum, L., et al., Different Patterns of Attention Modulation in Early N140 and Late P300 sERPs Following Ipsilateral vs. Contralateral Stimulation at the Fingers and Cheeks. Front Hum Neurosci, 2021. 15: p. 781778.

65. Matsuda, Y., et al., Event-Related Brain Potentials N140 and P300 during Somatosensory Go/NoGo Tasks Are Modulated by Movement Preparation. Brain Sci, 2023. 14(1).

66. Nakajima, Y. and N. Imamura, Relationships between attention effects and intensity effects on the cognitive N140 and P300 components of somatosensory ERPs. Clin Neurophysiol, 2000. 111(10): p. 1711–8.

67. Giersiepen, M., S. Schutz-Bosbach, and J. Kaiser, Freedom of choice boosts midfrontal theta power during affective feedback processing of goal-directed actions. Biol Psychol, 2023. 183: p. 108659.

68. McFerren, A., et al., Causal role of frontal-midline theta in cognitive effort: a pilot study. J Neurophysiol, 2021. 126(4): p. 1221–1233.

69. Du, Y.K., et al., Frontal-midline theta and posterior alpha oscillations index early processing of spatial representations during active navigation. Cortex, 2023. 169: p. 65–80.

70. Li, Y., et al., Global synchronization in the theta band during mental imagery of navigation in humans. Neurosci Res, 2009. 65(1): p. 44–52.

71. Thornberry, C., M. Caffrey, and S. Commins, Theta oscillatory power decreases in humans are associated with spatial learning in a virtual water maze task. Eur J Neurosci, 2023. 58(11): p. 4341–4356.

72. Heinrichs-Graham, E., et al., The functional role of post-movement beta oscillations in motor termination. Brain Struct Funct, 2017. 222(7): p. 3075–3086.

73. Jurkiewicz, M.T., et al., Post-movement beta rebound is generated in motor cortex: evidence from neuromagnetic recordings. Neuroimage, 2006. 32(3): p. 1281–9.

74. Pfurtscheller, G., K. Zalaudek, and C. Neuper, Event-related beta synchronization after wrist, finger and thumb movement. Electroencephalogr Clin Neurophysiol, 1998. 109(2): p. 154–60.

75. Zhang, X., et al., Beta rebound reduces subsequent movement preparation time by modulating of GABAA inhibition. Cereb Cortex, 2024. 34(2).

76. Guipponi, O., et al., fMRI Cortical Correlates of Spontaneous Eye Blinks in the Nonhuman Primate. Cereb Cortex, 2015. 25(9): p. 2333–45.

77. Eckstein, M.K., et al., Beyond eye gaze: What else can eyetracking reveal about cognition and cognitive development? Dev Cogn Neurosci, 2017. 25: p. 69–91.

78. Hanning, N.M., M.M. Himmelberg, and M. Carrasco, Presaccadic Attention Depends on Eye Movement Direction and Is Related to V1 Cortical Magnification. J Neurosci, 2024. 44(12).

79. Koevoet, D., et al., The Costs of Paying Overt and Covert Attention Assessed With Pupillometry. Psychol Sci, 2023. 34(8): p. 887–898.

80. Bozzacchi, C., et al., Awareness affects motor planning for goal-oriented actions. Biol Psychol, 2012. 89(2): p. 503–14.

81. Di Russo, F., et al., Beyond the “Bereitschaftspotential”: Action preparation behind cognitive functions. Neurosci Biobehav Rev, 2017. 78: p. 57–81.

82. Freude, G. and P. Ullsperger, Changes in Bereitschaftspotential during fatiguing and non-fatiguing hand movements. Eur J Appl Physiol Occup Physiol, 1987. 56(1): p. 105–8.

83. Bozzacchi, C., R.L. Cimmino, and F. Di Russo, The temporal coupling effect: Preparation and execution of bimanual reaching movements. Biol Psychol, 2017. 123: p. 302–309.

84. Haken, H., J.A. Kelso, and H. Bunz, A theoretical model of phase transitions in human hand movements. Biol Cybern, 1985. 51(5): p. 347–56.

85. Boecker, H., et al., A role of the basal ganglia and midbrain nuclei for initiation of motor sequences. Neuroimage, 2008. 39(3): p. 1356–69.

86. Errante, A., et al., *Activation of Cerebellum,* Basal Ganglia and Thalamus During Observation and Execution of Mouth, hand, and foot Actions. Brain Topogr, 2023. 36(4): p. 476–499.

87. Lehericy, S., et al., Motor control in basal ganglia circuits using fMRI and brain atlas approaches. Cereb Cortex, 2006. 16(2): p. 149–61.

88. Taniwaki, T., et al., Reappraisal of the motor role of basal ganglia: a functional magnetic resonance image study. J Neurosci, 2003. 23(8): p. 3432–8.

89. Ali, Y., V. Montani, and P. Cesari, Neural underpinnings of the interplay between actual touch and action imagination in social contexts. Front Hum Neurosci, 2023. 17: p. 1274299.

90. Verleger, R., et al., On how the motor cortices resolve an inter-hemispheric response conflict: an event-related EEG potential-guided TMS study of the flankers task. Eur J Neurosci, 2009. 30(2): p. 318–26.

91. Wang, H., Zheng, H., Wu, H., and Long, J., Behavior-Dependent Corticocortical Contributions to Imagined Grasping: A BCI-Triggered TMS Study. IEEE Transactions on Neural Systems and Rehabilitation Engineering, 2023. 31: p. 519–529.

92. Zaghi, S., et al., Inhibition of motor cortex excitability with 15Hz transcranial alternating current stimulation (tACS). Neurosci Lett, 2010. 479(3): p. 211–4.

93. Little, S. and P. Brown, The functional role of beta oscillations in Parkinson’s disease. Parkinsonism Relat Disord, 2014. 20 **Suppl 1**: p. S44–8.

94. Miocinovic, S., et al., Patterns of Cortical Synchronization in Isolated Dystonia Compared With Parkinson Disease. JAMA Neurol, 2015. 72(11): p. 1244–51.

95. Singh, A., et al., Mid-frontal theta activity is diminished during cognitive control in Parkinson’s disease. Neuropsychologia, 2018. 117: p. 113–122.

96. Bizovicar, N., et al., Decreased movement-related beta desynchronization and impaired post-movement beta rebound in amyotrophic lateral sclerosis. Clin Neurophysiol, 2014. 125(8): p. 1689–99.

